# A Novel Niclosamide Sulfate Prodrug with Enhanced Bioavailability Suppresses Hepatocellular Carcinoma via Inhibition of Multiple Signaling Pathways

**DOI:** 10.64898/2026.03.06.710248

**Authors:** Mingdian Tan, Steve Schow, Yi Liu, Robert Lum, Dawiyat Massoudi, Renumathy Dhanasekaran, Samuel So, Mei-Sze Chua

**Author notes:** Corresponding authors Mingdian Tan, Ph.D., Research Scientist, Department of Surgery, School of Medicine, Asian Liver Center, 1201 Welch Road, Stanford, Stanford, CA 94305, USA, Mei-Sze Chua, Ph.D., Senior Scientist, Department of Surgery, School of Medicine, Asian Liver Center, 1201 Welch Road, Stanford, Stanford, CA 94305, USA.

## Abstract

**Background:** Hepatocellular carcinoma (HCC) remains a leading cause of cancer-related mortality worldwide, underscoring the urgent need for effective therapies. The FDA-approved anthelmintic, niclosamide, is a promising repurposed drug candidate for HCC; however, its clinical application in solid tumors is hampered by poor aqueous solubility and resulting low bioavailability.

**Methods:** We designed and screened eight novel niclosamide prodrug candidates for solubility, stability, and anti-proliferative activity in HCC cell lines. The lead compound, SSL-0024, was evaluated for pharmacokinetics and anti-tumor efficacy in an orthotopic patient-derived xenograft (PDX) model, and for cytotoxicity in patient-derived organoids (PDOs). Mechanistic studies assessed AKT–mTOR–STAT3 and other related signaling pathways.

**Results:** The O-sulfate derivative, SSL-0024, demonstrated improved solubility and pH stability, and a sustained release of niclosamide over ∼48 h. Once-daily oral administration of SSL-0024 (100 mg/kg) achieved ∼60% tumor growth inhibition in PDX models at ∼46.8% of the niclosamide ethanolamine dose, with minimal toxicity. It also induced significantly greater cytotoxicity than sorafenib (*p<0.05*) in the PDO. Mechanistically, SSL-0024 suppressed major oncogenic pathways including AKT–mTOR–STAT3, RAF, and Wnt/β-catenin.

**Conclusions:** SSL-0024 overcomes key pharmacokinetic limitations of niclosamide while maintaining potent anti-tumor activity, supporting its further development as an orally bioavailable therapeutic candidate for HCC.

## Introduction

Hepatocellular carcinoma (HCC) is the most common type of primary liver cancer, representing approximately 75–90% of cases, ranking among the fifth to sixth most common cancers and the third leading cause of cancer-related mortality worldwide, with over 800,000 deaths annually^1^. Most patients present with advanced disease in the context of chronic liver injury. Despite recent advances in systemic therapy, sustained treatment responses remain limited due to molecular heterogeneity and compensatory activation of parallel oncogenic pathways^2^. Therapeutic strategies capable of simultaneously suppressing convergent survival networks while maintaining tolerable systemic exposure are urgently required.

Niclosamide, an FDA-approved anthelmintic drug, has emerged as a candidate for oncologic repurposing owing to its multi-target activity. Mechanistic studies have demonstrated that it suppresses multiple signaling pathways, including, but not limited to, Wnt/β-catenin signaling, through LRP6 degradation ^3^, STAT3-dependent transcriptional activity in cancer models^4^, and mTOR ^5^. Through coordinated interference with survival and proliferation pathways, niclosamide exhibits in vitro anti-tumor effects in HCC^5–7^ and other types of cancers ^8–13^.

However, despite robust preclinical efficacy, the clinical translation of niclosamide in solid tumors has been limited by extremely poor aqueous solubility and the resulting low oral bioavailability as well as rapid systemic clearance ^14^.

We previously reported that niclosamide effectively reversed HCC gene expression profiles to those of normal hepatocytes, suggesting its anti-HCC potential^5^. However, in vivo activity was only observed with a more water-soluble form, niclosamide ethanolamine (NEN), which is not FDA-approved for human use. To aid the clinical translation of niclosamide, we reported a proof-of-concept study using a water-soluble phosphate salt prodrug of niclosamide^6^, validating the well-accepted hypothesis^15^ that improving water solubility and consequent oral bioavailability is critical in enabling niclosamide to reach solid tumors to exert its multifaceted mechanisms. The phosphate salt approach is well established ^16^ and therefore lacks intellectual property potential; additionally, it causes a rapid spike in free niclosamide in the plasma, followed by rapid clearance, suggesting the need for frequent dosing and the potential for acute tissue toxicity with the sudden spike of niclosamide in tissues. Hoelzen et al. developed a valine niclosamide prodrug with enhanced water solubility and oral bioavailability; however, it lacked activity in a hollow-fiber assay of HCC cell line^17^. A stearate prodrug nanoparticle platform for niclosamide was also shown to enhance bioavailability after IV injection (not oral administration) in the treatment of osteosarcoma ^18, 19^. Given the limitations of the investigational niclosamide prodrugs, we aimed to design and develop novel niclosamide prodrugs that can enhance oral bioavailability and sustain longer systemic exposure, while preserving its previously validated antitumor mechanisms. We evaluated the lead compound, SSL-0024, for its pharmacokinetic profile, safety, and anti-tumor efficacy in orthotopic HCC patient-derived xenograft (PDX) models and confirmed its biological mechanisms of action.

## Materials and Methods

### Design, synthesis, and physicochemical characterization of novel niclosamide prodrug candidates

Based on existing patents and structure-activity relationship studies of niclosamide^20^, we designed over 50 novel potential prodrugs of niclosamide by modifying the 2-hydorxyl position with strongly anionic (SSL-0024, SSL-0049, and SSL-0053), neutral but polar (SSL-0048, SSL-0050, and SSL-0061), or lipophilic (SSL-0052 and SSL-0058) derivatives. Of these, eight were selected for synthesis by O2h Discovery (Cambridge, UK, and India) (Table 1). The chemical structures were confirmed by ^1^H-NMR spectroscopy, and the purity was determined by LC-MS/HPLC analysis (all chemicals were tested to have >95% purity). Detailed synthetic procedures and chemical analyses are provided in the Supplementary Materials.

**Table 1.**
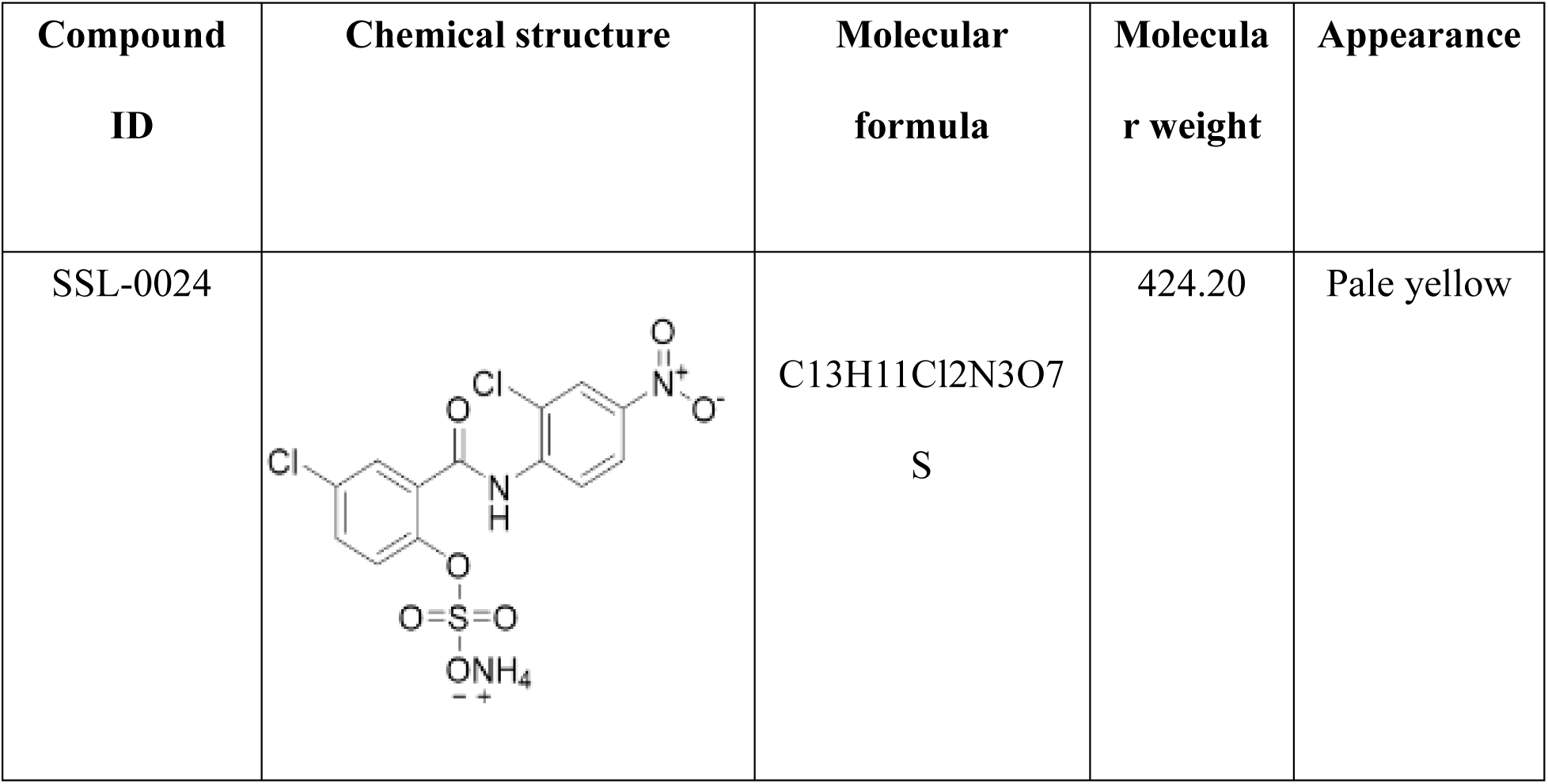
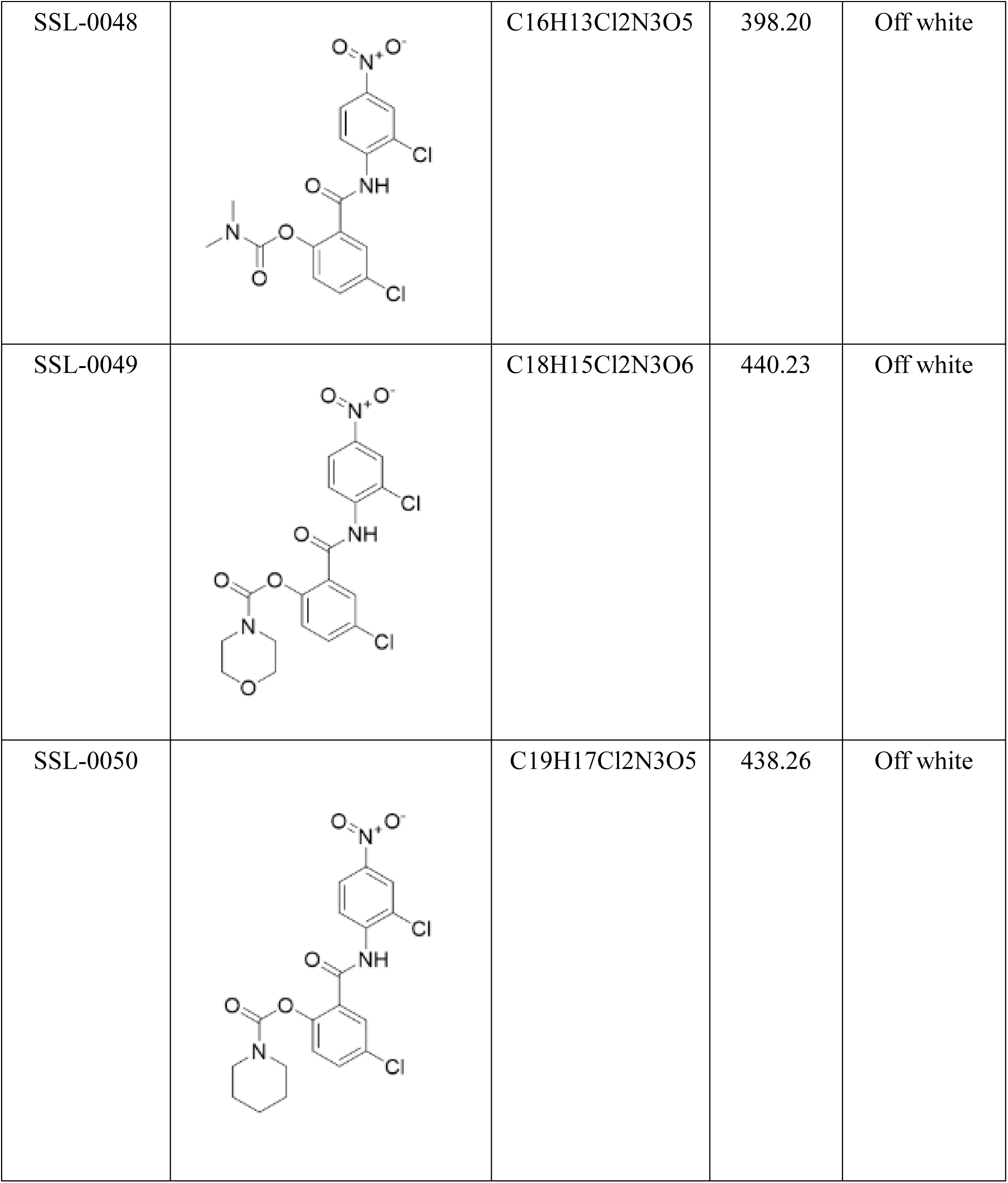
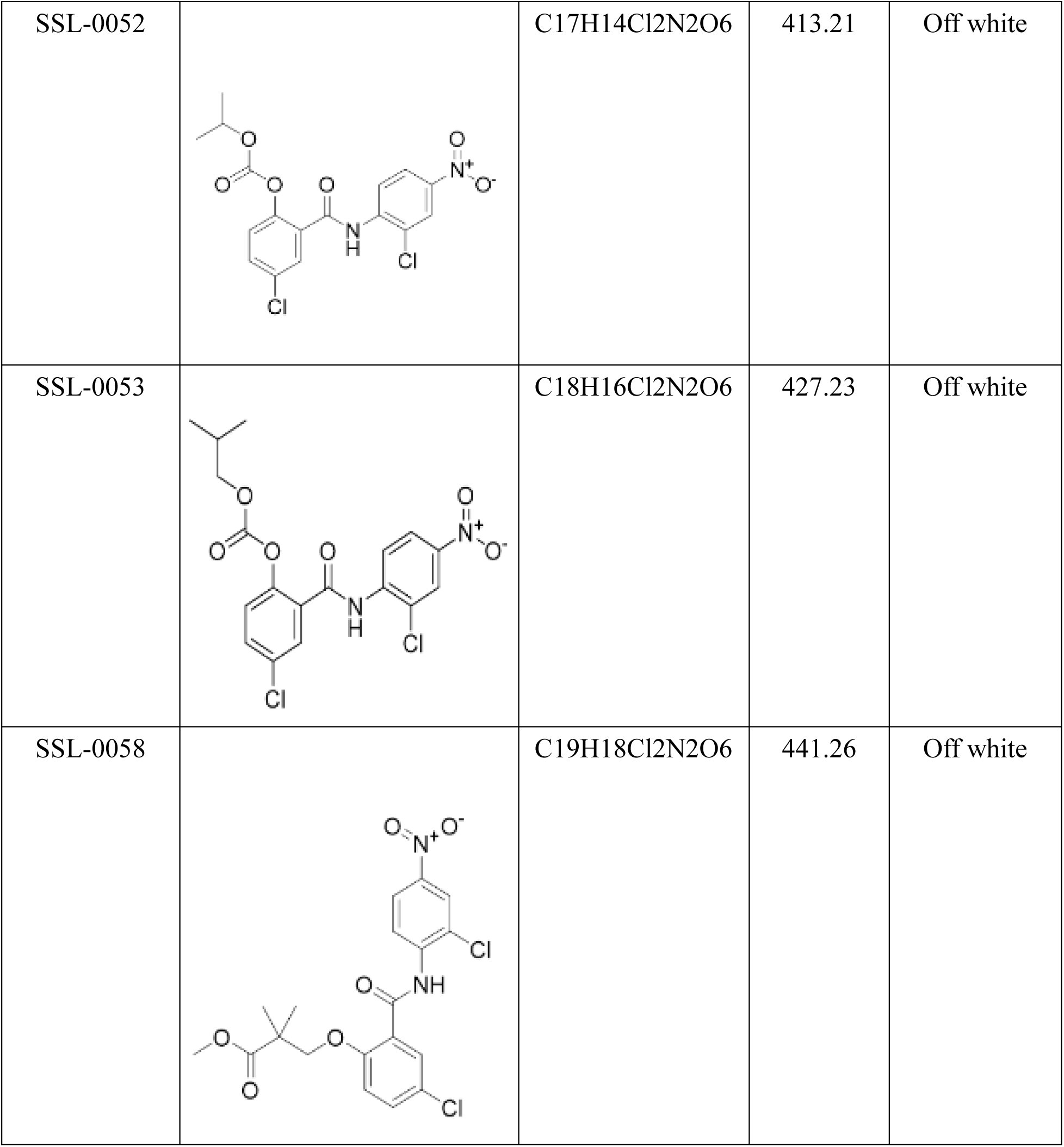
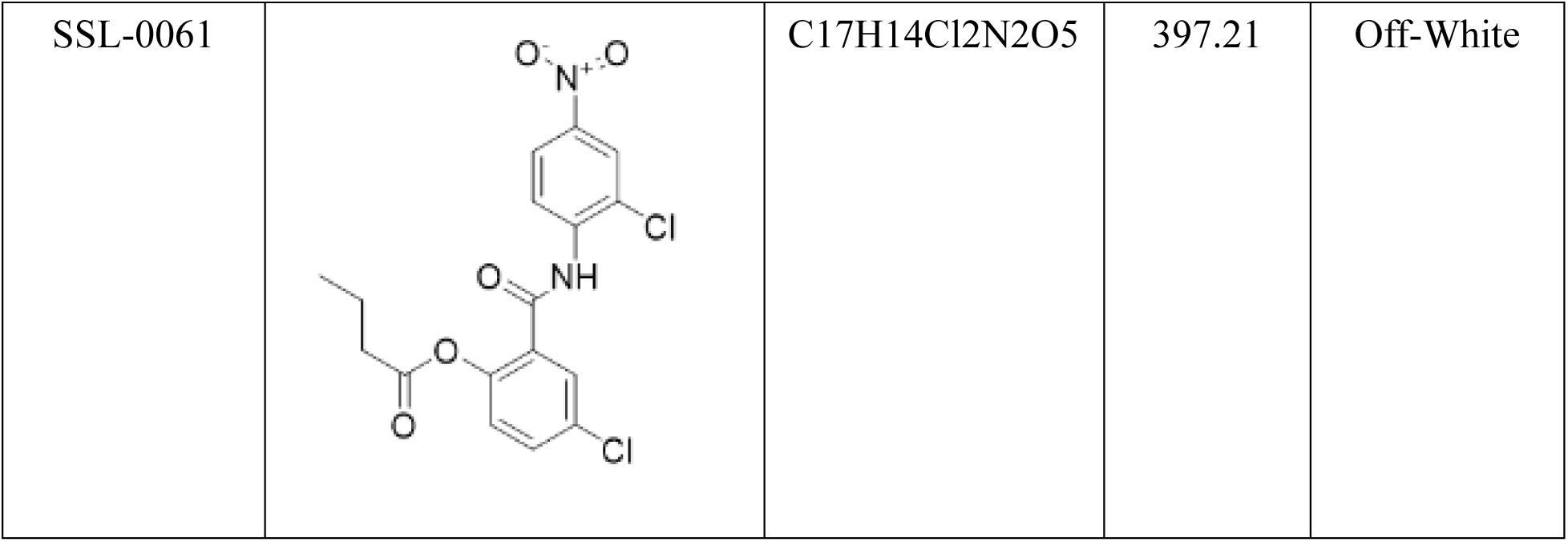
Compounds structure, molecular formula and molecular weight.

| **Compound ID** | **Chemical structure** | **Molecular formula** | **Molecular weight** | **Appearance** |
| --- | --- | --- | --- | --- |
| SSL-0024 |  | C13H11Cl2N3O7<br>S | 424.20 | Pale yellow |
| SSL-0048 |  | C<sub>16</sub>H<sub>13</sub>Cl<sub>2</sub>N<sub>3</sub>O<sub>5</sub> | 398.20 | Off white |
| SSL-0049 |  | C<sub>18</sub>H<sub>15</sub>Cl<sub>2</sub>N<sub>3</sub>O<sub>6</sub> | 440.23 | Off white |
| SSL-0050 |  | C<sub>19</sub>H<sub>17</sub>Cl<sub>2</sub>N<sub>3</sub>O<sub>5</sub> | 438.26 | Off white |
| SSL-0052 |  | C<sub>17</sub>H<sub>14</sub>Cl<sub>2</sub>N<sub>2</sub>O<sub>6</sub> | 413.21 | Off white |
| SSL-0053 |  | C<sub>18</sub>H<sub>16</sub>Cl<sub>2</sub>N<sub>2</sub>O<sub>6</sub> | 427.23 | Off white |
| SSL-0058 |  | C<sub>19</sub>H<sub>18</sub>Cl<sub>2</sub>N<sub>2</sub>O<sub>6</sub> | 441.26 | Off white |
| SSL-0061 |  | C17H14Cl2N2O5 | 397.21 | Off-White |

All eight prodrug candidates were assayed for their kinetic and thermodynamic water solubility, pH stability (2.0, 7.4), and in human plasma using standard assay protocols by O2h Discovery (Cambridge, India). Niclosamide (cat# N3510-50G, Millipore) and NEN (cat# 2A-9011637, 2A Biotech) were used as the reference controls.

### Culture of HCC cell lines

The human hepatocellular carcinoma (HCC) cell lines HepG2 (ATCC® HB-8065™, RRID: CVCL_0027) and Hep3B (ATCC® HB-8064™, RRID: CVCL_0326) were obtained from the American Type Culture Collection (ATCC), whereas Huh7 cells (JCRB0403, RRID: CVCL_0336) were provided by Dr. Mark Kay (Stanford University). HepG2 and Hep3B cells were cultured in Eagle’s Minimum Essential Medium (EMEM; ATCC, cat. no. 30-2003), and Huh7 cells were maintained in Dulbecco’s Modified Eagle’s medium (DMEM; ATCC, cat. no. 30-2002). All media were supplemented with 10% fetal bovine serum (FBS), 100 U/mL penicillin, and 100 μg/mL streptomycin (Life Technologies). Cells were maintained at 37°C in a humidified incubator with 5% CO₂.

### Cell proliferation assay

The anti-proliferative effects of the niclosamide prodrug candidates and niclosamide ethanolamine (used as a positive control) were assessed in HepG2, Huh7, and Hep3B cells using the CellTiter 96® Aqueous One Solution Cell Proliferation assay (MTS; Promega, cat. no. G4100). Cells were seeded into 96-well plates at a density of 3×10³ cells/well and allowed to adhere overnight. Each compound was used at a concentration range of 0.033–50 µM over an incubation period of 72 h. Following treatment, 15 μL of 3-(4,5-dimethylthiazol-2-yl)-5-(3-carboxymethoxyphenyl)-2-(4-sulfophenyl)-2H-tetrazolium inner salt (MTS) reagent was added to each well and incubated for 4 h at room temperature. The resulting formazan product was solubilized by adding 100 μL of Stop Solution per well and incubating at 37°C for 1 h before measuring the absorbance at 570 nm using a SYNERGY-LX microplate reader (BioTek). Dose–response curves and half-maximal inhibitory concentration (IC₅₀) values were determined by nonlinear regression analysis using the GraphPad Prism 11.0 (GraphPad Software, RRID:SCR_002798).

### Pharmacokinetic and Tissue Distribution Analysis in Orthotopic Patient-Derived Xenograft model of HCC

All animal experiments were conducted in accordance with the ARRIVE guidelines and approved by the Stanford University Institutional Animal Care and Use Committee (IACUC; Protocol APLAC-20167) in compliance with the USDA Animal Welfare Act, PHS Policy, and AAALAC International standards. Tumor size was monitored using calipers and did not exceed the maximum allowable diameter of 1.70 cm, with total tumor volume (calculated as W²×L/2) not exceeding 10% of the pre-inoculation body weight. Human HCC tissues were obtained from patients undergoing liver resection under an institutional review board-approved protocol, with informed consent. Six- to 10-week-old male NOD-scid IL2Rgamma^null^ mice (NSG mice, Transgenic, Knockout and Tumor Model Center, Stanford University) were housed under controlled environmental conditions (12-h light/dark cycle, 20–24°C, 40–60% humidity) with ad libitum access to food and water.

To provide a clinically relevant assessment of tumor and liver tissue exposure to niclosamide following administration of our prodrug candidates, we used an orthotopic HCC patient-derived xenograft (PDX) model, established using previously reported procedures^6^. Briefly, tumor fragments derived from resected patient samples were labeled with the luciferase reporter gene and surgically implanted into the livers of NSG mice after initial subcutaneous engraftment on the shoulders of NSG mice. Tumor formation was monitored by bioluminescence imaging following the injection of luciferin substrate (150 mg/kg) and monitored by Lago-X (Spectral Instruments Imaging).

Following confirmed tumor engraftment, PDX-bearing mice were administered a single dose of SSL-0024 (50 mg/kg) or niclosamide (40 mg/kg). At predetermined time points (5 min, 30 min, 1, 4, 8, 24, and 48 h post-administration), the animals were anesthetized with isoflurane, and serial blood samples (50 µL each) were collected via saphenous vein (thigh) puncture. Blood samples were immediately transferred into EDTA-coated microtubes, centrifuged to obtain plasma, and stored at −80°C until further analysis. Since SSL-0024 was expected to release free niclosamide in vivo, we assessed the levels of niclosamide in the plasma and harvested liver and PDX tissues using a validated LC–MS/MS method performed by Medicilon (Boston).

### In vivo efficacy study in orthotopic PDX HCC models

Following the procedure described above, PDX-bearing mice were randomized into three treatment groups (n=10 per group): vehicle control, SSL-0024 (100 mg/kg; molecular weight equivalent to 77.13 mg of niclosamide), or NEN (200 mg/kg, molecular weight equivalent to 168.57 mg of niclosamide). The compounds were administered once daily via oral gavage for 25 days.

Tumor progression was monitored weekly using bioluminescence imaging, and body weight was recorded three times per week. At study termination, PDX and adjacent liver tissues were harvested for niclosamide quantification using LC-MS/MS analysis (Medicilon, Boston) and immunohistochemical analysis. Blood was collected for analysis to determine its effects on blood biochemistry, liver and kidney functions, and lipid metabolism at the termination point (Stanford Animal Pathology Service).

Perioperative analgesia included Ethiqa-XR (3.25 mg/kg) administered prior to surgery, followed by postoperative carprofen (10–25 mg/kg) as needed. The animals were monitored regularly for tumor burden and general health. Humane endpoints were applied in accordance with the IACUC and AVMA guidelines. Euthanasia was performed using CO₂ inhalation or isoflurane anesthesia followed by cervical dislocation.

Tumor volume was quantified using standard volumetric calculations using the formula below:

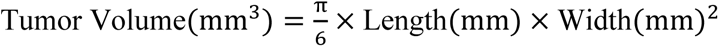

### Immunohistochemistry

Formalin-fixed, paraffin-embedded tissue sections were deparaffinized in xylene and rehydrated in graded ethanol to water solutions. Antigen retrieval was performed using citrate buffer (cat# k035; Diagnostic Biosystems) under heat-mediated conditions, followed by cooling to room temperature. Endogenous peroxidase activity was quenched using blocking buffer, and the sections were washed with phosphate-buffered saline (PBS).

Immunostaining was performed for the proliferation marker Ki-67 and the apoptotic marker cleaved caspase-3, following the manufacturer’s instructions for primary monoclonal antibodies against Ki-67 (1:20 dilution) (MA5-14520, Invitrogen^TM^, Waltham), and polyclonal antibodies against caspase-3 (1:300 dilution) (700182, Invitrogen). Briefly, slides were incubated with primary antibodies overnight at 4°C, followed by incubation with horseradish peroxidase (HRP)-conjugated secondary antibodies. Immunoreactivity was visualized using 3,3′-diaminobenzidine (DAB) substrate, and nuclei were counterstained with hematoxylin. After dehydration through graded ethanol and clearing in xylene, the slides were mounted for microscopic evaluation.

### Patient-derived organoids

Patient-derived organoids (PDOs) were established from human liver cancer tissue obtained from treatment-naive HCC patients undergoing resection under an institutional review board-approved protocol. The tissue was digested with DNase and Collagenase IV, treated with Erythrocyte Lysis Buffer (Qiagen), and the cells were seeded in a round bottom ultra-low attachment plate (7007, Costar) with organoid medium at 37°C in 5% CO₂ for 2-3 weeks until PDOs were formed. For drug treatment, PDOs were transferred to a 96 well plate and treated with DMSO (vehicle), sorafenib (10 µM), SSL-0024 (1 µM), or SSL-0024 (2 µM) for 48 h (n = 3 wells per condition). Viability was assessed using the Cyto3D Live-Dead Assay Kit (BM01, TheWell BIOSCIENCE), where Acridine Orange (AO) labeled live cells (green) and Propidium Iodide (PI) labeled dead cells (red). The spheroids were imaged at 10× magnification using a Keyence microscope. Image analysis was performed in R: a spheroid mask was generated from the brightfield channel using Gaussian smoothing and intensity thresholding, and the fluorescence signal was quantified within the masked region as integrated intensity above the background (median of non-spheroid pixels + 3 SD). Cell death was expressed as the dead/live ratio (PI integrated intensity/AO integrated intensity), and statistical comparisons between DMSO and each treatment group were performed using the t-test.

### Protein extraction and Western blotting

To assess the effects of niclosamide prodrug candidates on multiple signaling pathways, we treated Huh7 cells with a 10x × IC_50_ dose of SSL-0024 for 24 h, and protein levels were determined by western blotting.

Total protein was extracted from treated cells using T-PER Tissue Protein Extraction Reagent (Pierce, cat. no. 78510) and quantified using a BCA Protein Assay (Pierce, cat. no. 23225). Equal amounts of protein (20 μg) were resolved on 4–12% polyacrylamide gels (Invitrogen) and transferred to polyvinylidene difluoride (PVDF) membranes. The membranes were blocked in 5% nonfat milk in TBST for 1 h at room temperature and incubated overnight at 4°C with primary antibodies against LRP6 (Cell Signaling Technology, cat. no. 2560, RRID: AB_2139329, 1:1000 dilution), β-catenin (Cell Signaling Technology; cat. no. 8480, RRID: AB_11127855, 1:1000 dilution), STAT3 (Cell Signaling Technology, cat. no. 4904, RRID: AB_331269, 1:1000 dilution), phospho-STAT3 (Tyr705) (Cell Signaling Technology, cat. no. 9145, RRID: AB_2491009, 1:1000 dilution), Bcl-2 (Cell Signaling Technology, cat. no. 2870, RRID: AB_2290373, 1:1000 dilution), Bcl-xL (Cell Signaling Technology; cat. no. 2764, RRID: AB_2228008, 1:1000 dilution), and Cyclin D1 (Cell Signaling Technology, cat. no. 55506, RRID: AB_2827374, 1:1000 dilution), AKT (Cell Signaling Technology; cat. no. 9272S, RRID: AB_329827, 1:1000 dilution), phospho-AKT (Ser473) (Cell Signaling Technology, cat. no. 9271S, RRID: AB_329825, 1:1000 dilution), mTOR (Cell Signaling Technology; cat. no. 2972, RRID: AB_2105622, 1:1000 dilution), phospho-mTOR (Ser2448) (Cell Signaling Technology, cat. no. 2971, RRID: AB_10691552, 1:1000 dilution), phospho-p70 S6 Kinase (Thr389) (Cell Signaling Technology, cat. no. 9234, RRID: AB_2269803, 1:1000 dilution), phospho-4EBP1 (Thr37/46) (Cell Signaling Technology, cat. no. 2855, RRID: AB_560835, 1:1000 dilution), (epidmal growth factor receptor EGFR (; Cell Signaling Technology, cat. no. 2234, RRID: AB_331701, 1:1000 dilution), and A-Raf (Cell Signaling Technology, cat. no. 2752, RRID: AB_330813, 1:1000 dilution), and B-Raf (Cell Signaling Technology, cat. no. 9433, RRID: AB_2259354, 1:1000 dilution), and phospho-B-Raf (Ser445) (Cell Signaling Technology, cat. no. 2696, RRID: AB_390721, 1:1000 dilution), and C-Raf (Cell Signaling Technology, cat. no. 9422, RRID: AB_390808, 1:1000 dilution), and phospho-C-Raf (Ser338) (Cell Signaling Technology, cat. no. 9427, RRID: AB_2067317, 1:1000 dilution), and vasorin (Abcam, cat. no. ab156868, 1:500 dilution), PD-L1 (Cell Signaling Technology, cat. no. 13684, RRID: AB_2687655, 1:1000 dilution), ADAM17 (ABclonal, Cat. no. A0821, RRID: AB_2757412, 1:1000 dilution) and GAPDH (Proteintech, cat. no. 60004-1-Ig, RRID: AB_2107436, 1:20,000 dilution).

After washing, the membranes were incubated with fluorescent secondary antibodies (LI-COR Biosciences, IRDYE 800CW goat anti-rabbit, cat. no. 926-32211, RRID: AB_621843, 1:20,000 dilution; IRDYE 680RD goat anti-mouse, cat. no. 926-68070, RRID: AB_10956588, 1:20,000 dilution) at room temperature for 1 h. Signals were detected and recorded using an Odyssey imaging system (LI-COR Biosciences), according to the manufacturer’s instructions.

### Statistical analysis

Data are presented as mean ± standard error of the mean (SEM). Statistical analyses were performed using the GraphPad Prism software (version 11.0; GraphPad Software, RRID:SCR_002798). For comparisons among multiple groups, one-way analysis of variance (ANOVA) was applied, followed by Tukey’s post-hoc test when overall significance was achieved and variance homogeneity assumptions were satisfied.

To reduce inter-experimental variability, data were normalized to matched baseline or vehicle control values (expressed as fold change or percentage of control), where appropriate. Parametric analyses were conducted only after verification of normality using the Shapiro–Wilk test (α = 0.05) in conjunction with Q–Q plot assessment. For non-Gaussian distributions, log transformation was applied when normality improved; if assumptions remained violated, nonparametric tests (e.g., Mann–Whitney U test) were used. Nonparametric approaches were also employed when normalized datasets exhibited zero variance within a reference group (e.g., control SEM = 0). Statistical significance was defined as a two-tailed P-value < 0.05.

## Results

### Niclosamide prodrug design strategy

Because of its poor solubility (1.6 mg/L) and high melting point (230°C °C), niclosamide is minimally absorbed from the gastrointestinal tract, with very low recovery of the drug or its metabolites from blood or urine^21^. To make niclosamide more bioavailable and clinically translatable to address HCC, a small exploratory collection of niclosamide phenolic derivatives was prepared as potential solubilizing prodrugs. The niclosamide phenol group was modified with highly charged functional groups for water solubilization, neutral but polar functional groups for water solubilization without a charge, and lipophilic derivatives for aiding in membrane passage and lowering the melting point (Table 1). The strongly anionic derivatives included O-sulfate (SSL-0024), 3-sulfopropanic ester (SSL-0049), and 4-phosphate-2-methylbutanoic ester (SSL-0053); the neutral but polar derivatives included 2-(dimethyl amino) acetic ester (SSL-0048), O-sulfonamide (SSL-0050), and 4-(glucoside)-oxy-butanoic ester (SSL-0061); and the lipophilic derivatives included O-methylene pivalate (SSL-0052) and 4-(isobutyryloxy) butanoic ester (SSL-0058).

### Physicochemical and in vitro activity profiling and lead prodrug selection

The above candidates were systematically evaluated for kinetic and thermodynamic solubility, chemical stability, plasma stability, and in vitro antiproliferative activity to identify the lead compound(s) suitable for in vivo development (Table 2). Niclosamide and NEN were used as reference controls.

**Table 2.** Physicochemical properties and in vitro potency of niclosamide prodrug candidates.

| Compound | Kinetic solubility | Thermodynamic solubility | | pH7.4 (2h) | | pH2.0 (2 h) | | Human Plasma (2 h) | | IC50 (μM) |
| --- | --- | --- | --- | --- | --- | --- | --- | --- | --- | --- |
| | μM | μM | Equivalent conc. in μg/mL | % Remaining | % Niclosamide | % Remaining | % Niclosamide | % Remaining | % Niclosamide | (Average in 3 cell lines) |
| **SSL-0024** | **15** | **2** | **1** | **88** | **0** | **94** | **1.1** | **85.6** | **0.1** | **3.87** |
| SSL-0048 | 0.5 | 1 | 0 | 21.8 | 12.7 | 11.4 | 2.4 | 93.4 | 0.3 | NA |
| SSL-0049 | 0.1 | 0.60 | 0.2 | 29.8 | 0.7 | 22.8 | 0.6 | 75.9 | 18.3 | NA |
| SSL-0050 | 0.4 | 0.7 | 0.3 | 64.9 | 21.5 | 2.1 | 0.7 | 91.9 | 0.8 | NA |
| **SSL-0052** | **3** | **3.0** | **1.3** | **76.2** | **22.3** | **37.4** | **32** | **0** | **71.9** | **2.51** |
| **SSL-0053** | **0.3** | **0.2** | **0.1** | **0** | **77.8** | **23.7** | **41.6** | **0** | **68.8** | **2.6** |
| SSL-0058 | 1.3 | 3.8 | 1.7 | 36 | 57.9 | 13.4 | 39.5 | 92.8 | 1.1 | NA |
| SSL-0061 | 0.6 | 0.4 | 0.2 | 54.50 | 30.80 | 9.40 | 6.30 | 0.0 | 59.4 | 1.445 |
| Niclosamide | 6.9 | 0.5 | 0.2 | 85.2 | NA | 9.6 | NA | 96.8 | NA | 1.02 |
| NEN | 2.7 | 3.6 | 1.4 | 105.3 | NA | 11.9 | NA | 83.2 | NA | 1.03 |

The lipophilic derivatives (SSL-0052 and SSL-0058) displayed enhanced thermodynamic solubility relative to niclosamide; this most likely resulted from the lipid functionalities disrupting the crystal packing features of planar aromatic niclosamide. The O-sulfate derivative SSL-0024 showed significantly enhanced kinetic (2.2-fold increase) and thermodynamic (5-fold increase) solubility compared to niclosamide. Unexpectedly, the other two strongly charged derivatives (SSL-0049 and SSL-0053) had solubilities comparable to or lower than those observed for niclosamide. Neutral but polar derivatives (SSL-0048, SSL-0050, and SSL-0061) containing both lipophilic and polar components did not enhance the solubility of niclosamide.

Additionally, SSL-0024 demonstrated high chemical stability and resistance to pH-driven decomposition under both physiological (pH 7.4) and gastric (pH 2.0) conditions, with minimal conversion to the parent compound over a 2-hour incubation period, while the other derivatives showed varying degrees of hydrolytic instability under both pH conditions. In human plasma, 85.6% of SSL-0024 remained intact, indicating limited premature hydrolysis and suggesting the potential for controlled systemic release. As for human plasma stability, only three derivatives (SSL-0052 - lipophilic, SSL-0053 - anionic, and SSL-0061 – neutral/polar) displayed significant instability. These derivatives also exerted anti-proliferative activity in HCC cells, which could be attributed to the liberation of niclosamide, which then directly acted on the cells. The remaining derivates were stable in plasma; of these, only SSL-0024 showed anti-proliferative activity in HCC cell lines (mean IC₅₀ = 3.87 µM), Other derivatives (SSL-0048, 0049, 0050, 0058)failed to show anti-proliferative effects presumably because they do not cross the cell membrane or do not release niclosamide once inside the cell.

Collectively, these data identified SSL-0024 as a lead prodrug candidate for further pharmacokinetic and in vivo efficacy evaluation based on its favorable solubility, stability, and biological activity profiles.

Kinetic and thermodynamic solubilities were measured under standardized assay conditions. Chemical stability and conversion to niclosamide were assessed following a 2-hour incubation at pH 7.4, pH 2.0, and in human plasma. Percentage remaining indicates an intact prodrug, and percentage niclosamide reflects conversion to the parent compound. IC₅₀ values represent average antiproliferative activity across three cancer cell lines. Niclosamide and NEN were included as reference controls, based on previous reports. NA: not available.

### SSL-0024 Exhibits Extended Pharmacokinetics and Favorable Tissue Distribution

Plasma concentration–time profiling revealed a marked extension of systemic exposure to SSL-0024 when administered as a single oral dose of 50 mg/kg to NSG mice bearing orthotopic HCC PDX. Specifically, SSL-0024 resulted in a prolonged release of free niclosamide in the plasma over 48 h (Fig. 1A). Niclosamide reached maximal plasma concentrations at 5 min (Tmax = 5 min), followed by a rapid elimination phase characterized by a sharp decline in systemic exposure. The same PK profile was observed with our proof-of-concept niclosamide phosphate salt prodrug ^6^. Additionally, area under the plasma concentration–time curve from 0 to 24 h (AUC₀–₂₄h) showed that SSL-0024 exhibited 6.6 times greater exposure than niclosamide within the shared 24-h observation window (Fig. 1B), demonstrating greatly enhanced oral bioavailability compared to niclosamide.

**Figure 1.**
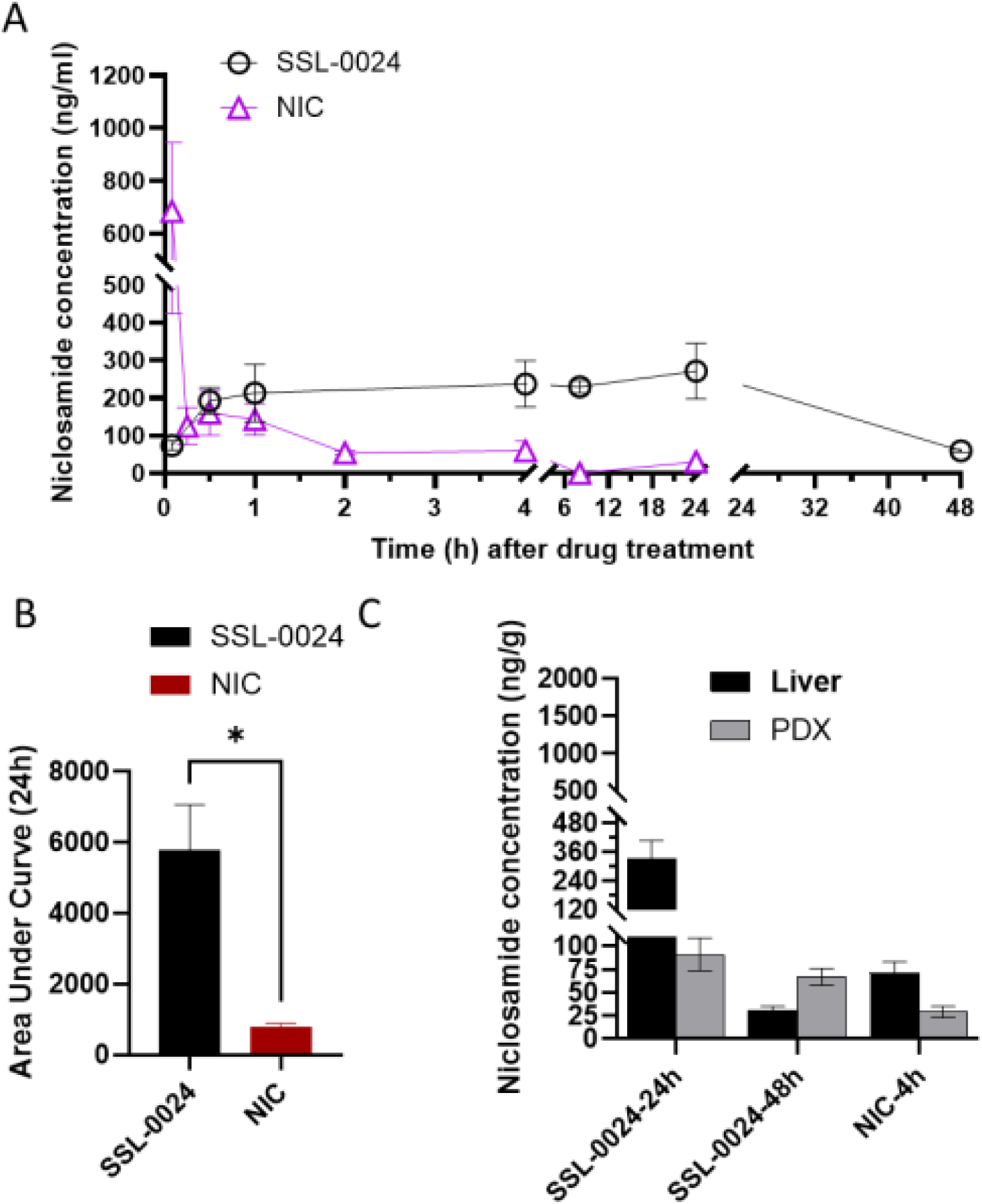
Pharmacokinetic and tissue distribution analysis of Niclosamide prodrugs. (A) Plasma concentration–time profiles of niclosamide following oral administration of SSL-0024 or niclosamide (both compounds formulated in 0.5% methylcellulose). (B) Area under the plasma concentration–time curve from 0–24 h (AUC₀–₂₄h). Oral administration of SSL-0024 resulted in 6.6 times greater plasma exposure of niclosamide than when NIC was administered within the shared 24 h observation window. (C) Quantification of released niclosamide levels in liver and orthotopic PDX tumor tissues following administration of SSL-0024 or niclosamide. Plasma and tissue concentrations were quantified by LC-MS/MS. Data are presented as mean ± SEM.

After 24 or 48 h, we sacrificed the mice and harvested liver and PDX tissues to assess the amount of free niclosamide released from SSL-0024. Tissue distribution analysis revealed marked differences in niclosamide exposure between liver and PDX tumors. At 24 h following SSL-0024 administration, liver niclosamide concentrations were substantially higher than those in the PDX tissue (332.6 vs 91.4 ng/g). In contrast, at 48 h, niclosamide preferentially accumulated in PDX tumors compared to liver (66.8 vs 31 ng/g) (Fig. 1C). In comparison, NIC treatment resulted in consistently higher niclosamide accumulation in the liver than in the PDX tumors at the indicated time points (Fig. 1C). The remaining candidates exhibited a limited tissue exposure (Fig. S2). These findings suggest that SSL-0024 provides a substantially improved pharmacokinetic profile with sustained plasma and tissue exposure.

Collectively, these results demonstrated that SSL-0024 achieves prolonged systemic exposure, enhanced overall drug exposure (significantly increased AUC), and favorable in vivo conversion to niclosamide. These pharmacokinetic advantages support its selection as a lead candidate for subsequent efficacy evaluations in orthotopic HCC models.

### SSL-0024 Suppresses *in vivo* Growth of Orthotopic Patient-derived Xenograft of HCC

For the in vivo efficacy study, PDX-bearing mice were treated with either 100 mg/kg SSL-0024 (molecular weight equivalent to 77.13 mg of niclosamide) or 200 mg/kg NEN (molecular weight equivalent to 168.57 mg of niclosamide) via once-daily oral gavage for four weeks. NEN was used as the reference control rather than niclosamide, since we had previously shown that niclosamide lacks in vivo activity ^5^. The 100 mg/kg dose of SSL-0024 was selected based on our previous study of the niclosamide phosphate salt prodrug ^6^), which demonstrated strong tumor inhibition at a comparable molecular equivalent. SSL-0024 reduced the tumor volume by 59.5% relative to the control, whereas NEN reduced the tumor volume by 43.7% (Fig. 2A–B). Although the difference between SSL-0024 and NEN was not statistically significant, SSL-0024 showed a trend toward greater tumor growth inhibition. Notably, this effect was achieved with a lower molecular weight equivalent of niclosamide (approximately 45.8% of that in the NEN group).

**Figure 2.**
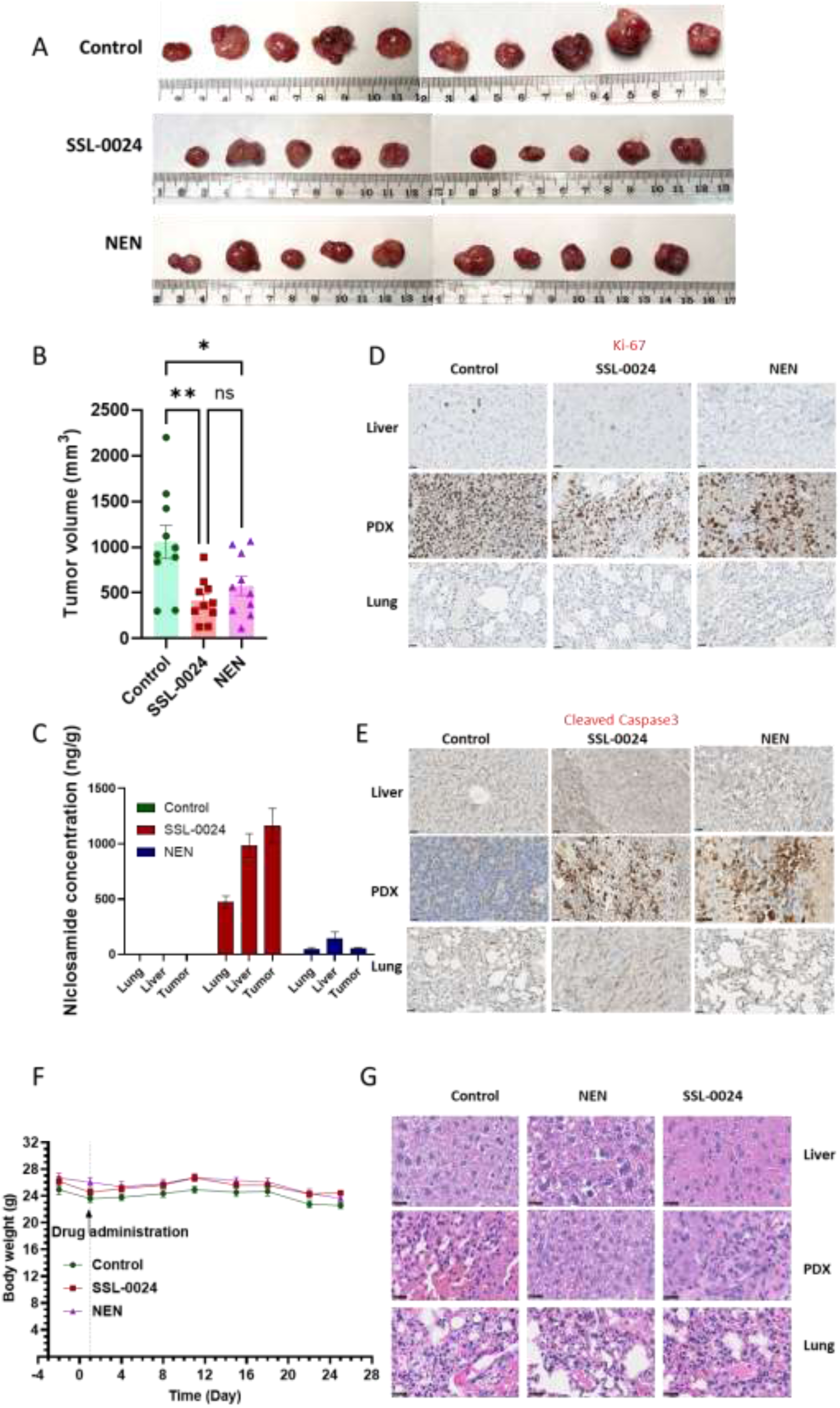
SSL-0024 suppresses HCC PDX tumor growth and modulates proliferation and apoptosis markers. (A) Images of all harvested tumors collected at the end of the 25-day oral treatment period with SSL-0024 (100 mg/kg) or NEN (200 mg/kg). (B) Tumor volume measurements and statistical analysis over the 25-day treatment period (n = 10 per group). SSL-0024 reduced tumor volume by 59.5% compared to control, while NEN reduced tumor volume by 43.7%. Although the difference between SSL-0024 and NEN was not statistically significant, SSL-0024 showed a trend toward greater tumor inhibition. (C) Niclosamide concentrations in PDX tumor, liver, and lung tissues following treatment (n = 5 per group), quantified by LC-MS/MS. SSL-0024 achieved markedly higher accumulation in liver (981.79 vs. 142.97 ng/g) and tumor (1,161.89 vs. 53.89 ng/g) tissues compared to NEN, with preferential tumor distribution. Data are presented as mean ± SEM. (D–E) Immunohistochemical staining of Ki-67 (D) and cleaved caspase-3 (E) in PDX tumors. SSL-0024 demonstrated stronger inhibition of proliferation (Ki-67) and similar induction of apoptosis (cleaved caspase-3) relative to NEN; magnification: ×80. (F) Body weight of tumor-bearing mice monitored throughout the 25-day treatment period. No significant body weight loss was observed across treatment groups, suggesting that SSL-0024 and NEN were well tolerated at the doses administered. Data are presented as mean ± SEM (n = 10). (G) Representative hematoxylin and eosin (H&E) staining of liver, PDX tumor, and lung tissues. No obvious histopathological abnormalities were observed in liver or lung tissues following treatment, indicating acceptable systemic toxicity profiles for both SSL-0024 and NEN; magnification: ×80.

Analysis of niclosamide concentrations in PDX tumors and the liver revealed significantly higher drug accumulation with SSL-0024 than with NEN (Fig. 2C). While NEN-treated mice exhibited ∼1.7-fold higher niclosamide levels in the normal adjacent liver versus PDX tissue, SSL-0024-treated mice showed ∼1.2-fold higher niclosamide levels in PDX tumors than in the normal liver (Fig. 2C), indicating preferential tumor accumulation. Additionally, niclosamide levels in normal livers after SSL-0024 administration were approximately 6-fold higher than those achieved with NEN, further supporting its enhanced bioavailability and tissue distribution. We also assessed the distribution of niclosamide in the lung tissues to determine lung toxicity. Niclosamide exposure in the lungs was markedly lower than that in the liver, corresponding to 0.48 the liver levels after SSL-0024 administration and 0.34 times after NEN treatment (Fig. 2C). At the molecular level, SSL-0024 exhibited stronger inhibition of Ki-67 expression than NEN, suggesting enhanced anti-proliferative activity, whereas both compounds induced comparable increases in cleaved caspase-3, indicating similar pro-apoptotic effects (Fig. 2D–E). Collectively, these results demonstrate that SSL-0024 not only improves bioavailability and tumor-targeted accumulation, but also achieves potent tumor growth inhibition with a favorable safety profile, highlighting its strong translational potential.

Body weight measurements throughout the treatment period revealed no significant weight loss in the SSL-0024 treated group compared to that in the control and NEN-treated groups (Fig 2F), suggesting its overall safety. Previous reports have suggested that NEN at 200 mg/kg q.d. induces alveolar toxicity due to degradation products, including ethanolamine^22^; consistent with this, we observed similar pathological changes in the lungs induced by NEN, but not in SSL-0024 treated group (Fig. 2G). Biochemical analysis of blood parameters, liver and kidney functions, and lipid metabolism indicated that treatment with SSL-0024 significantly increased WBC, neutrophil counts, alkaline phosphatase, and altered lipid profiles (cholesterol, HDL, LDL) compared to the control, without major changes in RBC parameters or the liver enzyme AST (ALT showed elevated levels with high variability but remained within the normal range). NEN treatment had modest effects on MCV, MCH, monocyte %, AST, and alkaline phosphatase levels. Both compounds maintained normal RBC morphology and overall hematologic homeostasis, indicating tolerable systemic effects at the doses tested (Table S2).

### SSL-0024 inhibited growth of patient-derived organoids

To determine whether PDOs can be an alternative, reliable HCC model for evaluating SSL-0024 and other novel compounds, we treated PDO derived from one HCC patient with DMSO (vehicle), sorafenib at 10 µM, or SSL-0024 at 1 µM or 2 µM for 48 h, and assessed PDO viability by AO/PI live/dead staining. DMSO-treated organoids maintained a high live/dead ratio with predominantly green (AO+) fluorescence, confirming spheroid integrity. Sorafenib treatment resulted in a modest but non-significant increase in cell death compared to DMSO treatment (p = 0.089) (Fig. 3A-3E). In contrast, SSL-0024 induced a dose-dependent increase in cell death, with both 1 µM (p = 0.025) and 2 µM (p = 0.007) doses significantly elevating the PI/AO ratio above the DMSO control, indicating that the majority of cells within treated spheroids were non-viable. SSL-0024 also significantly increased cell death compared to sorafenib at both 1 µM (p = 0.042) and 2 µM (p = 0.022), suggesting its greater potency than the standard-of-care agent.

**Figure 3.**
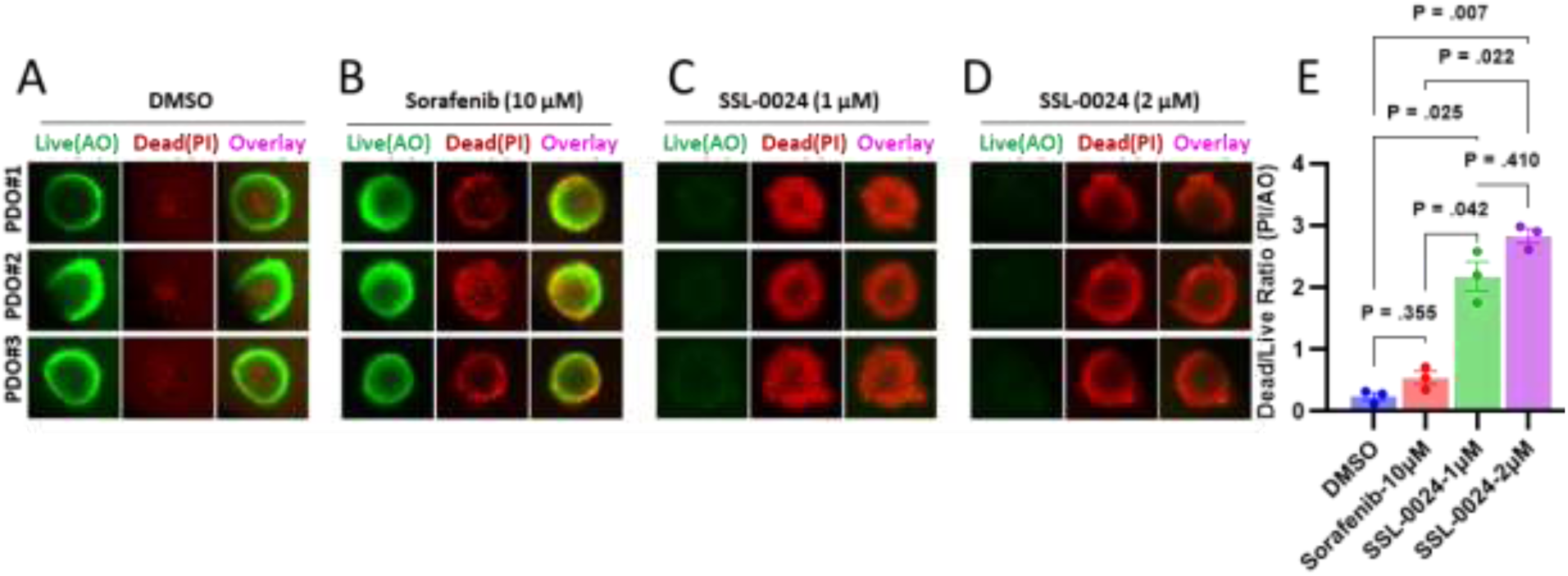
SSL-0024 induces a dose-dependent increase in cell death in PDOs. Fluorescence images of PDOs stained with AO/PI live/dead staining after a 48-hour treatment with (A) DMSO, (B) 10 µM of sorafenib, (C)1 µM of SSL-0024, or (D) or 2 µM of SSL-0024. Image analysis shows a modest increase in cell death in the PDOs treated with sorafenib (B) while the PDOs treated with SSL-0024 present a significant dose-dependent increase in cell death (C and D) compared to the DMSO treated PDOs (A). (E) is the quantification of the fluorescence images shown in (A), (B), (C) and (D).

### SSL-0024 regulates HCC growth *via* suppressing AKT–mTOR–STAT3, RAF, and Wnt signaling

To elucidate the mechanism underlying SSL-0024–mediated HCC growth inhibition, we focused on the AKT–mTOR–STAT3 signaling axis, a key pathway previously reported to be regulated by niclosamide ^5^. Protein levels were normalized to the control (set as 1), and relative changes were assessed for the SSL-0024 and NEN groups.

SSL-0024 treatment significantly suppressed the phosphorylation of critical signaling proteins, including p-STAT3 (Y705, 0.51), p-AKT (0.037), p-mTOR (0.28), p-4EBP1 (0.085), and p-p70S6K (0.18) (Fig. 4A-D). Total STAT3 and total AKT were also significantly reduced by SSL-0024, whereas mTOR showed no statistically significant difference compared to the control (Fig. 4A-D), indicating that SSL-0024 primarily inhibits the active, phosphorylated forms of these proteins. In contrast, NEN treatment led to a relatively modest reduction in STAT3, whereas BCL-XL was significantly decreased in the NEN group but remained unchanged in SSL-0024–treated tumors (Fig. 3A-D). Cyclin D1, a key downstream regulator of proliferation, was significantly reduced by both SSL-0024 and NEN (Fig. 4A and 4H).

**Fig. 4.**
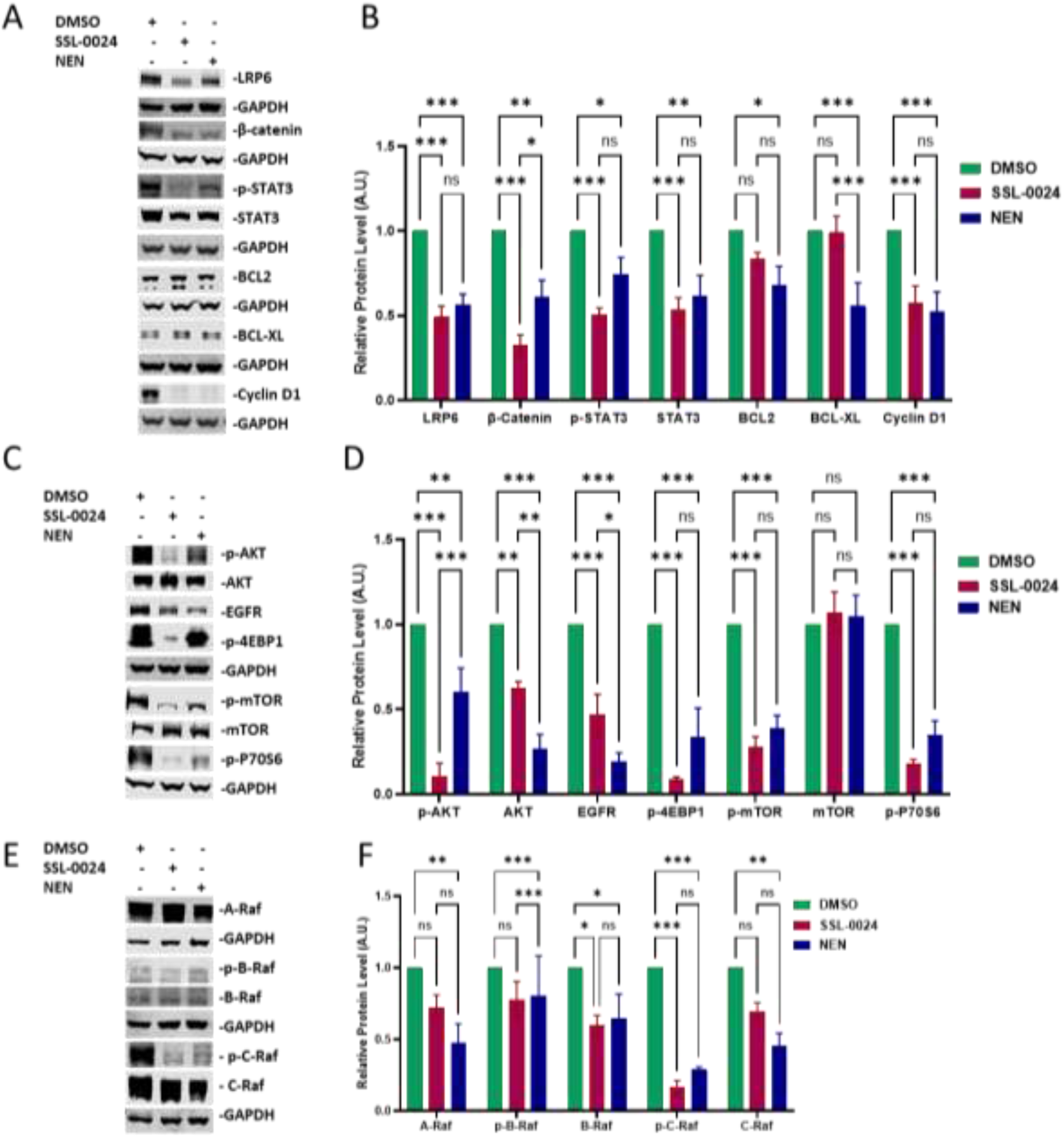
SSL-0024 modulates multiple signaling pathways. (A) Representative immunoblot analysis of LRP, β-catenin, p-STAT3 (Y705), total STAT3, BCL2, BCL-XL, and Cyclin D1 protein levels following SSL-0024 treatment. (B) Quantification of protein expression levels shown in (A). (C) Representative immunoblot analysis of p-AKT, total AKT, EGFR, p-4EBP1, p-mTOR, total mTOR, and p-p70S6K following SSL-0024 treatment. (D) Quantification of protein expression levels shown in (C). (E) Representative immunoblot analysis of total A-Raf, p-B-Raf, total B-Raf, p-C-Raf, and total C-Raf following SSL-0024 treatment. (F) Quantification of protein expression levels shown in (E). Protein levels were quantified using densitometry and expressed as arbitrary units (A.U.). Phosphorylated proteins were normalized to their respective total protein levels, except p-4EBP1, which was normalized to GAPDH due to the unavailability of a reliable total 4EBP1 signal. All other proteins were normalized to GAPDH. Data are presented as the relative protein levels (A.U. ± SEM). n = 3. *p < 0.05, **p < 0.01, ***p < 0.001 vs. control.

SSL-0024 treatment significantly reduced the expression of total A-Raf, total B-Raf, total C-Raf, and phosphorylated C-Raf, whereas phosphorylated B-Raf showed no statistically significant changes (Fig. 4E–F; p-A-Raf could not be detected). These results indicate that SSL-0024 effectively suppresses RAF signaling, primarily through the inhibition of C-Raf activation, a key driver of upstream MAPK signaling in HCC^23^. In contrast, NEN treatment showed no statistically significant changes in A-Raf, B-Raf, or phosphorylated B-Raf, while other proteins, including C-Raf and phosphorylated C-Raf, were significantly decreased. The expression of EGFR, which is an activator of both the AKT and RAS signaling cascades ^24, 25^, was significantly reduced by both SSL-0024 and NEN (Fig. 4A-D).

Two key components of the Wnt signaling pathway, LRP6 and β-catenin, were significantly downregulated by both SSL-0024 and NEN (Fig. 4A-D).

Overall, these results demonstrate that SSL-0024 suppresses HCC growth through the inhibition of multiple crucial signaling pathways, and it achieves stronger inhibitory effects on most of the phosphorylated proteins than NEN, correlating with its enhanced in vivo anti-tumor effects.

### SSL-0024 regulates HCC growth *via* suppressing VASN and PD-L1 protein expression

Consistent with our previous study using niclosamide phosphate salt ^6^, SSL-0024 strongly inhibited the expression of VASN and PD-L1 (Fig. 5A and 5 B). To assess the upstream regulation of VASN, we measured ADAM17, the metalloprotease responsible for VASN cleavage, and the subsequent activation of TGFβ signaling ^26^. SSL-0024 treatment decreased ADAM17 levels (Fig. 5A and 5 B), suggesting reduced VASN secretion and suppression of TGFβ-mediated protumor signaling.

**Fig. 5.**
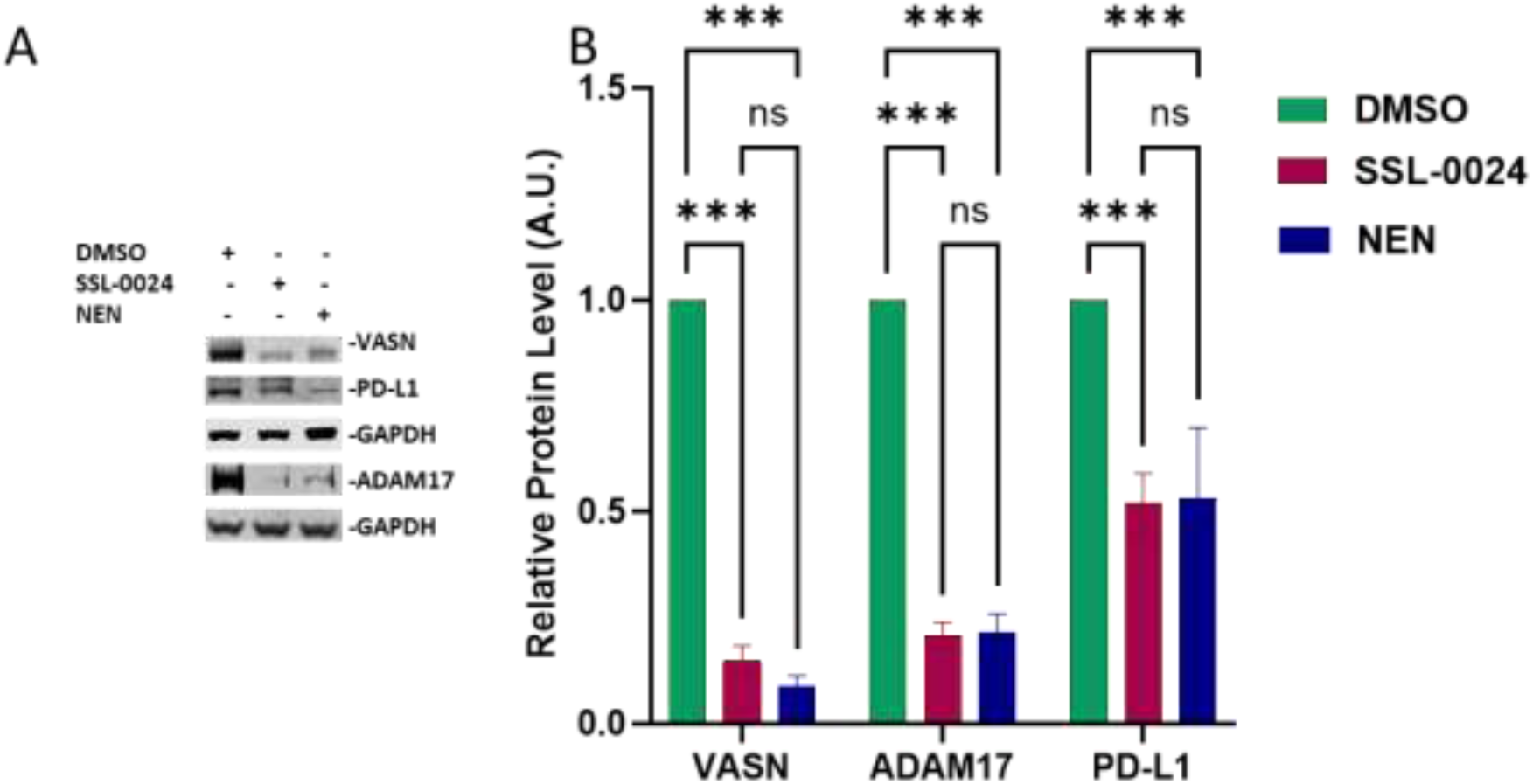
SSL-0024 regulates VASN-associated protein expression. (A) Representative immunoblot analysis of VASN, PD-L1, and ADAM17 following SSL-0024 treatment. (B) Quantification of protein expression levels shown in (A). Protein levels were quantified using densitometry and expressed as arbitrary units (A.U.). All the proteins were normalized to GAPDH. Data are presented as the relative protein levels (A.U. ± SEM). n = 3. \**p* < 0.05, \*\**p* < 0.01, \*\*\**p* < 0.001 vs. control.

In addition to the validated changes in oncogenic signaling pathways, we showed that SSL-0024 acts via the novel target VASN and its associated proteins, as well as PD-L1, supporting its multi-targeted mechanism and superior anti-tumor potential.

## Discussion

By rationally designing and systematically screening eight novel prodrug candidates of niclosamide, we identified a lead prodrug, SSL-0024 (niclosamide-O-sulfate ammonium salt), which showed favorable water solubility, pH stability, improved PK profile/oral bioavailability, and in vitro and in vivo anti-tumor efficacy using multiple HCC models. In particular, SSL-0024 achieves sustained systemic exposure beyond 24 h after a single dose, whereas native niclosamide exhibits a plasma half-life averaging 4–5 h ^27^. This prolonged half-life enhances bioavailability, so that there is a steady level of active parent drug reaching the tumors; consequently, we observed effective tumor suppression at only ∼45.8% of the dose required for NEN, without detectable systemic toxicity. By reducing the dosing requirement while maintaining efficacy, SSL-0024 demonstrates a substantially improved therapeutic index, supporting once-daily oral administration and highlighting its potential for clinical use in patients with HCC. This pharmacokinetic improvement directly addresses the historical barrier that has limited the clinical translation of niclosamide and potentially enables sustained inhibition of multiple oncogenic pathways critical for HCC growth and survival.

Phenolic sulfates typically have a poor reputation as prodrugs because of their general stability (which our study corroborated). The strongly charged O-sulfate also strongly inhibits passive membrane passage. Therefore, our observation that the O-sulfate prodrug SSL-0024 demonstrated enhanced oral bioavailability and preservation of the biological activity of niclosamide in vitro and in vivo was unexpected. Additionally, niclosamide is known to be rapidly cleared from the bloodstream, as observed in our study and others, which may be because its phenolic group is a target for sulfotransferases to facilitate water absorption and clearance via the kidneys ^28^. In contrast, SSL-0024 remained in the bloodstream for over 24 h, with a steady release of niclosamide over 48 h, as well as the accumulation of free niclosamide in the liver and PDX tissues, which presents a clinical advantage by increasing tumor exposure and reducing dosing frequency.

Although O-sulfates are excellent water-solubilizing moieties, they require the intervention of a sulfatase enzyme to liberate the bioactive phenolic drug. In our study, aryl sulfatases in liver cells^29^ were expected to cleave SSL-0024 to release free niclosamide. The ammonium sulfate counterion that is also released upon cleavage is labeled by the Food and Drug Administration as being “Generally Recognized As Safe” (GRAS) and is a commonly used food additive safe for general human consumption. Having a safe counterion is an important criterion in the selection of counterions in prodrug design^30, 31^ and is what we have achieved in this study. In contrast, NEN dissociates into free niclosamide and ethanolamine, which is a known lung irritant ^32^, and is likely responsible for the pathological changes in lung alveoli that we observed upon NEN treatment. Lung toxicity observed upon NEN administration was not observed following SSL-0024 treatment. Therefore, SSL-0024 is an unexpected niclosamide prodrug that is more efficacious and less toxic than NEN and has not been approved by the FDA for human use.

In our PDO HCC model, SSL-0024 exhibited potent cytotoxic activity in a dose-dependent manner, with its efficacy exceeding that of sorafenib under the same treatment conditions. The partial effect of sorafenib is consistent with prior literature^33^, and positions SSL-0024 as a potentially more effective alternative in HCC therapy. Importantly, the PDO HCC model preserves the three-dimensional architecture and cellular heterogeneity of primary tumors, features that are lost in conventional monolayer cultures and are known to influence drug penetration and metabolism. Therefore, PDOs represent a cost-effective, time-efficient, and clinically relevant screening platform uniquely suited to evaluating not only cytotoxic efficacy but also safety, mechanism of action, and combination therapy strategies for novel drug candidates such as SSL-0024. The ability of SSL-0024 to overcome the inherent diffusion barriers and drug resistance associated with three-dimensional growth further strengthens its translational potential.

Mechanistically, we confirmed that SSL-0024 suppresses a convergent oncogenic signaling network central to HCC progression, consistent with the reported actions of its parent drug, niclosamide. We observed coordinated inhibition of the mTOR–AKT–STAT3 axis, as evidenced by reduced phosphorylation of AKT, mTOR, 4EBP1, p70S6K, and STAT3 (Y705), together with downregulation of Cyclin D1, BCL2, and BCL-XL. Concurrent attenuation of RAF isoforms suggests disruption of growth factor–driven mitogenic signaling ^34^. By simultaneously inhibiting the mTOR–AKT–STAT3 and RAF–MEK–ERK cascades, SSL-0024 may prevent compensatory signaling that often drives resistance to monotherapies. This network-level suppression is clinically relevant, as multikinase inhibitors such as sorafenib rely on similar pathways, and resistance frequently arises via adaptive PI3K–AKT or RAF–MEK–ERK activation ^35–38^. Sustained STAT3 inhibition is particularly important given its established role in promoting proliferation, survival, and immune evasion in liver cancer ^39–41^. Simultaneous multi-pathway inhibition may limit the adaptive resistance often observed with single-target therapies, providing a rationale for more durable tumor control in HCC ^42^.

SSL-0024 reduced LRP6 expression to a degree comparable to that in NEN; however, β-catenin suppression was notably more pronounced following SSL-0024 treatment. Since β-catenin is a key downstream effector of LRP6-mediated Wnt signaling and drives the transcription of proliferation-associated genes^43^, the disproportionate reduction in β-catenin suggests that SSL-0024 may modulate β-catenin stability or activity through mechanisms beyond receptor-level suppression. These findings suggest that enhanced attenuation of canonical Wnt/β-catenin signaling may contribute, at least in part, to the superior anti-proliferative effects of SSL-0024.

In our previous study on niclosamide phosphate salt prodrug ^6^, VASN knockdown phenocopied drug treatment and VASN rescue attenuated antitumor effects, establishing VASN as a functional mediator of niclosamide-derived activity. Consistent with this mechanism, SSL-0024 reduced VASN expression, presumably via the release of free niclosamide. Together, these findings indicate that SSL-0024 conserves the previously validated VASN-dependent signaling axis and that our rational structural modification does not compromise mechanistic targeting, reinforcing confidence in its antitumor activity.

Beyond tumor-intrinsic signaling, SSL-0024 reduced PD-L1 expression, a central mediator of immune escape in HCC and a clinically actionable target ^44^. Although the precise regulatory mechanism underlying PD-L1 downregulation by SSL-0024 remains to be determined, one possible contributing mechanism might be the suppression of STAT3 signaling, a well-known transcriptional regulator of PD-L1 in multiple cancers such as lung cancer, natural killer/T-cell lymphoma, breast cancer, head and neck squamous cell carcinoma, and HCC^45–51^. This dual inhibition of the proliferative and immune-evasive pathways suggests that SSL-0024 could provide additive or synergistic effects when combined with immune checkpoint inhibitors.

Future studies should evaluate SSL-0024 in combination with immune checkpoint or kinase inhibitors for potential synergistic efficacy. Its ability to modulate both tumor growth and immune evasion highlights the potential for rational combination strategies with the current standard-of-care therapies.

From a translational perspective, SSL-0024 offers the additional advantage of simple one-step synthesis from readily available chemical precursors, facilitating scalable production and potentially reducing costs. Compared with current therapies, including Sorafenib, Lenvatinib, or checkpoint inhibitors such as Atezolizumab, SSL-0024 combines multipathway tumor suppression with potential immune modulation in a single orally available agent. Together with prolonged systemic exposure, reduced effective dosing, and minimal toxicity, these attributes position SSL-0024 as a pharmacologically optimized derivative with improved drug-like properties compared to earlier niclosamide formulations.

This study has several limitations that warrant consideration. First, although SSL-0024 was observed to suppress key signaling pathways, formal pharmacokinetic–pharmacodynamic correlation analyses were not performed, which would strengthen the mechanistic linkage between systemic exposure and target inhibition. Second, although PD-L1 reduction was observed, comprehensive characterization of the tumor immune microenvironment and assessment of immune cell infiltration were beyond the scope of this study. Third, the efficacy of the combination of approved HCC therapies, including sorafenib, lenvatinib, and immune checkpoint inhibitors, remains to be evaluated. Fourth, dose-escalation studies to define the maximum tolerated dose and therapeutic window, particularly in orthotopic PDX models, would provide valuable translational insights. Finally, additional investigations into other potentially relevant signaling networks, such as insulin-like growth factor regulation, may identify further opportunities for therapeutic synergy.

## Conclusion

In summary, SSL-0024 exemplifies the transformation of a biologically validated, but clinically impractical compound into a therapeutically viable drug candidate for solid tumors. Enhanced and sustained systemic exposure, together with coordinated oncogenic pathway suppression, suggests the promising potential of SSL-0024 as a readily translatable niclosamide prodrug for HCC therapy. Our findings lay a strong foundation for further mechanistic exploration, combination therapy studies, and translational development toward clinical applications, offering a compelling path forward for niclosamide-based therapeutic strategies in HCC.

## Abbreviations

¹H-NMR: proton nuclear magnetic resonance
ANOVA: analysis of variance
AUC: area under curve
DAB: 3,3′-diaminobenzidine
FBS: fetal bovine serum
GRAS: generally recognized as safe
HCC: hepatocellular carcinoma
HPLC: high-performance liquid chromatography
HRP: horseradish peroxidase
IACUC: Institutional Animal Care and Use Committee
IC₅₀: half-maximal inhibitory concentration
LC-MS: liquid chromatography-mass spectrometry
MTS: 3-(4,5-dimethylthiazol-2-yl)-5-(3-carboxymethoxyphenyl)-2-(4-sulfophenyl)-2H-tetrazolium, inner salt
NEN: niclosamide ethanolamine
NSG: NOD scid gamma
PDX: patient-derived xenografts
PVDF: polyvinylidene difluoride
SEM: standard error of the mean;

## Acknowledgement

We are grateful to the Stanford Cell Sciences Imaging Facility for providing training and usage support with RRIDSCR_017787.

## Author Contributions

Conceived study: Mei-Sze Chua, Steve Schow, Robert Lum, Samuel So. Design of compounds: Steve Schow and Robert Lum. Designed experiments: Mingdian Tan, Mei-Sze Chua and Renumathy Dhanasekaran. Participated in/executed trial/study: Mingdian Tan, Yi Liu, and Dawiyat Massoudi. Data collection and acquisition: Mingdian Tan, Yi Liu, and Dawiyat Massoudi. Data analysis: Mingdian Tan, Mei-Sze Chua, Dawiyat Massoudi. Data interpretation: Mingdian Tan, Steve Schow, Mei-Sze Chua, and Renumathy Dhanasekaran. Manuscript writing: Mingdian Tan, Mei-Sze Chua, and Steve Schow. Supervision of study: Mei-Sze Chua, Renumathy Dhanasekaran, Samuel So. Approval of final submitted version: Mingdian Tan, Mei-Sze Chua, Steve Schow, Robert Lum, Yi Liu, Dawiyat Massoudi, Renumathy Dhanasekaran, and Samuel So.

## Data availability

The datasets used and/or analyzed during the current study are available from the corresponding author upon reasonable request.

---

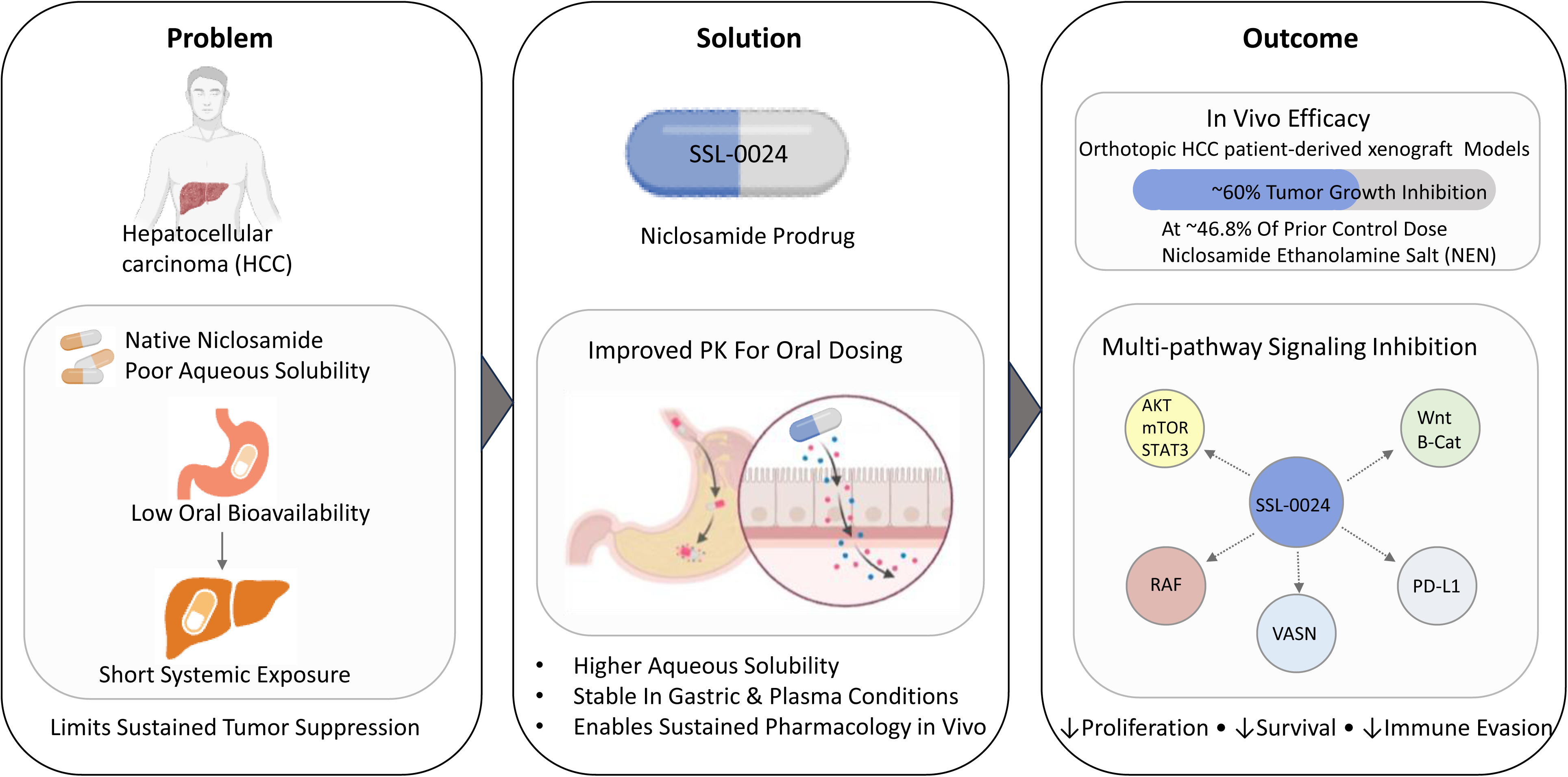

