## supplemental materials for "A Novel Niclosamide Sulfate Prodrug with Enhanced Bioavailability Suppresses Hepatocellular Carcinoma via Inhibition of Multiple Signaling Pathways"

**Abbreviations**

| HPLC | High-performance liquid chromatography |
| --- | --- |
| LCMS | Liquid Chromatograph-Mass Spectrometry |
| NMR | Nuclear Magnetic Resonance |
| TLC | Thin layer chromatography |
| Rt | Retention time |
| rt | Room temperature |
| °C | Degree Celsius |
| h | Hour/hours |
| ACN | Acetonitrile |
| DCM | Dichloromethane |
| EtOAc | Ethyl acetate |
| MeOH | Methanol |
| THF | Tetrahydrofuran |
| DMSO | Dimethyl sulfoxide |
| TEA | Triethylamine |
| DIPEA | N, N-Diisopropylethylamine |
| DMAP | 4-Dimethylaminopyridine |
| TPP | Triphenylphosphine |
| DIAD | Diisopropyl azodicarboxylate |
| Na_2_SO_4_ | Sodium sulfate |
| NaHCO_3_ | Sodium bicarbonate |
| HCl | Hydrochloric acid |

**1 ANALYTICAL DETAILS**

**NMR**: 1H-NMR spectra were recorded on Bruker AVANCE NEO 400MHz spectrometers in deuterated solvents. Chemical shifts (δ) are reported in parts per million and coupling constants (J values) in hertz. Spin multiplicities are indicated by the following symbols: s (singlet), d (doublet), t (triplet), q (quartet), m (multiplet), and bs (broad singlet). Deuterated solvents are given in parentheses and have a chemical shift of dimethyl sulfoxide (δ 2.50 ppm), methanol (δ 3.32 ppm), chloroform (δ 7.28 ppm), or other solvent as indicated in NMR spectral data.

**MS**: Mass Spectra were obtained on a Waters Acquity QDa spectrometer with empower and a water Acquity H-class equipped with a PDA spectrometer with empower software. Chromatography was performed using the C-18 column and suitable solvent as indicated in specific examples.

**Flash Column Chromatography System**: Flash purification was conducted with a Biotage Isolera with HP-Sil or KP-NH SNAP cartridges (Biotage) and Combi-*flash RF^+^* TELE DYNE ISCO. The solvent gradient is indicated in specific examples.

**Thin layer chromatography (TLC)**: TLC was carried out on silica gel plates with UV detection.

**LCMS Method**

| **Method Name** | **Column used** | **Method Description** |
| --- | --- | --- |
| o2h_LCMS_Method-A | X-Bridge BEH C18, 50 x 2.1 mm, 2.5 micron | Water Acquity UPLC- H Class equipped with PDA and attached with QDa detector, Column temperature: Ambient, Auto sampler temperature: 150C, Mobile Phase A : 2 mM ammonium acetate followed by 0.1%Formic acid in water, Mobile Phase B : 0.1% Formic Acid in Acetonitrile, Mobile phase gradient details: T = 0 min (95% A, 5% B) flow; T = 0.4 min (95% A, 5% B) ; gradient to T = 0.8 min (65% A, 35% B) ; gradient to T = 1.20 min (45% A, 55% B) ; T = 2.5 min (0% A, 100% B) ; gradient to T= 3.30 min (0% A, 100% B) ; gradient to T= 3.31 min to end of run at T = 4 min (95% A, 5% B), Flow rate: 0.55 mL/min, Run Time: 4 min. UV Detection Method: PDA Mass parameter: Probe: ESI, Mode of Ionisation : positive and negative, Cone voltage : 10V and 30V, capillary voltage: 0.8 KV, Extractor Voltage: 1KV, Rf Lens: 0.1,Temperature of source: 120°C,Temperature of Probe: 600°C, Cone Gas Flow:- Default , Desolvation Gas flow:-Default. |

**HPLC Method**

| **Method Name** | **Column used** | **Method Description** |
| --- | --- | --- |
| o2h_HPLC_Method_A | Waters Sunfire C18 (150mm x 4.6mm)3.5µm     or Equivalent | Machine Details: Water alliance e2695 with 2998 PDA detector Column temperature: 25°C Auto sampler temperature: 25°C,Mobile Phase A: 0.1% FA in Water Mobile Phase B: 100% Acetonitrile Mobile phase Gradient :- T = 0 min (90% A, 10% B) T = 7 min (10%A, 90% B)  T = 9 min (0% A, 100% B) T = 14 min (0% A, 100% B)  T = 14.01 min (90% A, 10% B) gradient to  T= 17 min (90% A, 10% B) Flow: 1mL/min Run Time: - 17 min UV Detection Method: - PDA |
| o2h_HPLC_Method_B | X-bridge C18 (150mm x 4.6mm) 3.5µm or Equivalent | Machine Details: Shimadzu 2050c with PDA detector Column temperature: 25°C Auto sampler temperature: 25°C,Mobile Phase A: 5mM Ammonium Bicarbonate in Water Mobile Phase B: 100% Acetonitrile Mobile phase Gradient:- T = 0 min (90% A, 10% B) T = 7 min (10%A, 90% B)  T = 9 min (0% A, 100% B) T = 14 min (0% A, 100% B)  T = 14.01 min (90% A, 10% B) gradient to  T= 17 min (90% A, 10% B) Flow: 1mL/min Run Time: - 17 min UV Detection Method: - PDA |

### 2. Synthetic procedures and chemical analyses

**Experimental of SSL-0024 (APX-X-0008)**

**Synthesis of Ammonium 4-chloro-2-((2-chloro-4-nitrophenyl) carbamoyl) phenyl sulfate**

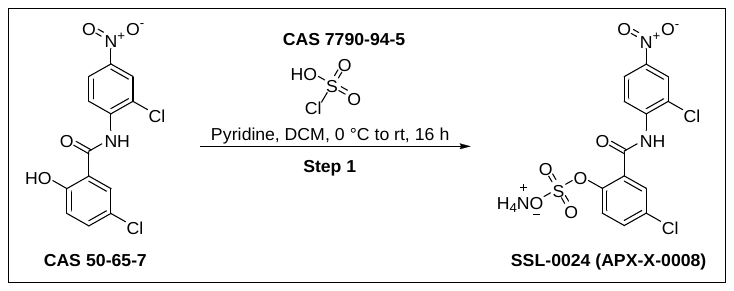

**Synthesis of** **Ammonium** **4-chloro-2-((2-chloro-4-nitrophenyl) carbamoyl) phenyl sulfate**

To a stirred solution of 5-chloro-N-(2-chloro-4-nitrophenyl)-2-hydroxybenzamide (CAS 50-65-7, 0.25 g, 0.766 mmol) and pyridine (5.0 mL) in DCM (2.0 mL) was added Chlorosulfonic acid (CAS 7790-94-5, 1.0 mL) dropwise at 0 °C and stirred at rt for 16 h. The reaction mixture was diluted with water (20 mL) and extracted with DCM (2 x 20 mL). The combined organic layer was washed with aqueous Citric acid solution (25 mL) and dried over Na_2_SO_4_, filtered and concentrated under reduced pressure. The crude material was purified by Prep HPLC to afford 4-chloro-2-((2-chloro-4-nitrophenyl) carbamoyl) phenyl sulfate as a pale-yellow solid (0.025 g, 0.059 mmol, 7% yield). ¹H NMR (400 MHz, DMSO-*d*_6_): *δ* 10.67 (s, 1H), 8.64 (d, *J* = 9.2 Hz, 1H), 8.42 (d, *J =* 2.8 Hz*,* 1H), 8.29 (dd, *J =*2.8, 2.8 Hz, 1H), 7.92 (q, *J* = 1.2 Hz, 1H), 7.66 (t, *J* = 1.2 Hz, 2H), 7.08 (brs, 3H), LCMS: O2h_LCMS_Method_A, Rt: 1.613 min, [M-H]^+^: 405.1.

**Experimental of SSL-0048 (APX-X-0010)**

**Synthesis of 4-chloro-2-((2-chloro-4-nitrophenyl) carbamoyl) phenyl dimethylcarbamate**

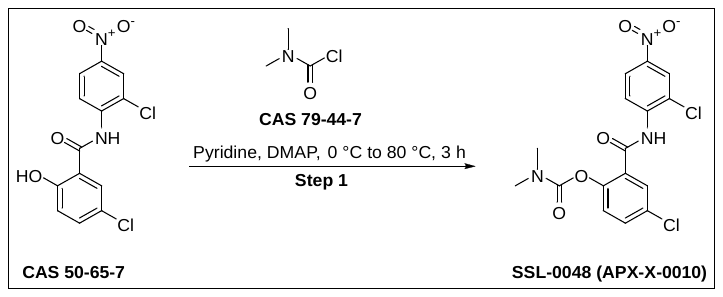

**Synthesis of 4-chloro-2-((2-chloro-4-nitrophenyl) carbamoyl) phenyl dimethylcarbamate**

To a stirred solution of 5-chloro-N-(2-chloro-4-nitrophenyl)-2-hydroxybenzamide (CAS 50-65-7; 0.3 g, 0.920 mmol) in pyridine (6.0 mL) were added DMAP (0.06 gm, 0.552 mmol) and dimethylcarbamic chloride (CAS 79-44-7, 0.227 g, 2.116 mmol) at 0 °C and heated at 80 °C for 3 h. The reaction mixture was diluted with aqueous Citric acid solution (50.0 mL) and extracted with EtOAc (2 x 50 mL). The combined organic layers were dried over Na_2_SO_4_, filtered and concentrated under reduced pressure. The crude material was purified by flash chromatography (Biotage, Normal phase, Silica gel, Mesh size - 230 - 400, 15% EtOAc in hexane) followed by washed with aqueous 1N HCl solution (25.0 mL; to remove traces of pyridine) to afford 4-chloro-2-((2-chloro-4-nitrophenyl) carbamoyl) phenyl dimethylcarbamate as an off-white solid (0.07 g, 0.17 mmol, 19% yield). ¹H NMR (400 MHz, DMSO-*d_6_*): δ 10.42 (s, 1H), 8.41 (d, *J* = 2.8 Hz, 1H), 8.28 (dd, *J =* 2.8, 2.8 Hz, 1H), 8.08 (d, *J* = 8.8 Hz, 1H), 7.77 (d, *J* = 2.4 Hz, 1H), 7.66 (dd, *J =* 2.8, 2.4 Hz, 1H), 7.35 (d, *J* = 8.8 Hz, 1H), 3.00 (s, 3H), 2.86 (s, 3H), LCMS: O2h_LCMS_Method_A, Rt: 2.324 min, [M-H]^+^: 396.2.

**Prep HPLC: MOA**

**Instrument Name: PHP-08-WATERS 2545 QUATERNARY SYSTEM WITH WATERS 2489 UV Detector**

Chromatographic separation and isolation were conducted with PHP-08-WATERS 2545 QUATERNARY SYSTEM WITH WATERS 2489 UV Detector. The column used was X-BRIDGE C18(150 x 19 mm ID, 5μm) and the compounds were eluted with**, Mobile Phase A**: 5MM AMMONIUM BICARBONATE + 0.05% AMMONIA IN WATER, **Mobile Phase B :** ACETONITRILE: WATER(80:20) with a gradient of T = 0.00 min (70% A, 30% B); gradient to T = 15.00 min (56% A, 44% B), T = 26.00 min (56% A, 44% B), T = 26.01 min (0% A, 100% B), T = 28.00 min (0% A, 100% B), T = 28.01 min (70% A, 30% B), T = 30.00 min (70% A, 30% B), Flow rate=13 mL/min; analysis time 30 min.

**Experimental of SSL-0049 (Target A)**

**Synthesis of 4-chloro-2-((2-chloro-4-nitrophenyl)carbamoyl)phenyl morpholine-4-carboxylate**

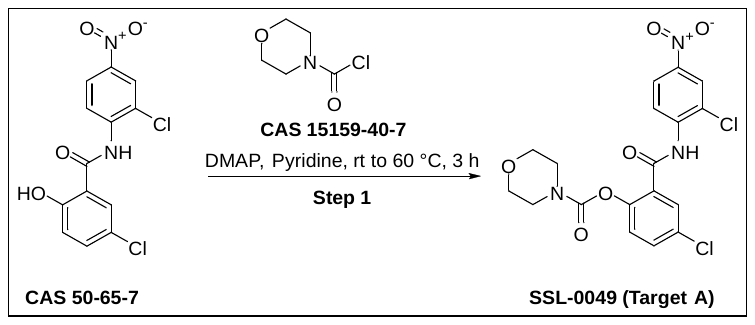

**Synthesis of 4-chloro-2-((2-chloro-4-nitrophenyl) carbamoyl) phenyl morpholine-4-carboxylate**
To a stirred solution of 5-chloro-N-(2-chloro-4-nitrophenyl)-2-hydroxybenzamide (CAS 50-65-7, 0.1 g, 0.30 mmol) in pyridine (2.0 mL) were added DMAP (CAS 1122-58-3, 0.022 g, 0.18 mmol) and piperidine-1-carbonyl chloride (CAS 15159-40-7, 0.105 g, 0.70 mmol) at rt and stirred at 60 °C for 3 h. The reaction mixture was diluted with water (30 mL) and extracted with EtOAc (3 x 15 mL). The combined organic layers were washed with 0.1N HCl solution (30 mL), dried over Na_2_SO_4_, filtered and concentrated under reduced pressure. The crude material was purified by flash chromatography (Biotage, Normal phase, 230 – 400 mesh silica, 18% EtOAc in n-hexane) to afford 4-chloro-2-((2-chloro-4-nitrophenyl) carbamoyl) phenyl morpholine-4-carboxylate as an off-white solid (0.075 g, 0.17 mmol, 74% yield). ¹H NMR (400 MHz, DMSO-*d*_6_): *δ* 10.46 (s, 1H), 8.42 (d, *J* = 2.8 Hz, 1H), 8.29 (dd, *J* = 2.8, 2.4 Hz, 1H), 8.07 (d, *J* = 9.2 Hz, 1H), 7.78 (d, *J* = 2.8 Hz ,1H), 7.68 (dd, *J* = 2.4, 2.4 Hz, 1H), 7.37 (d, *J* = 8.4 Hz, 1H), 3.54 (s, 6H), 3.34 (s, 2H); LCMS: O2h_LCMS_Method_A, Rt: 2.34 min, [M-H]^+^: 438.0.

**Experimental of SSL-0050 (Target B)**

**Synthesis of 4-chloro-2-((2-chloro-4-nitrophenyl) carbamoyl) phenyl piperidine-1-carboxylate**

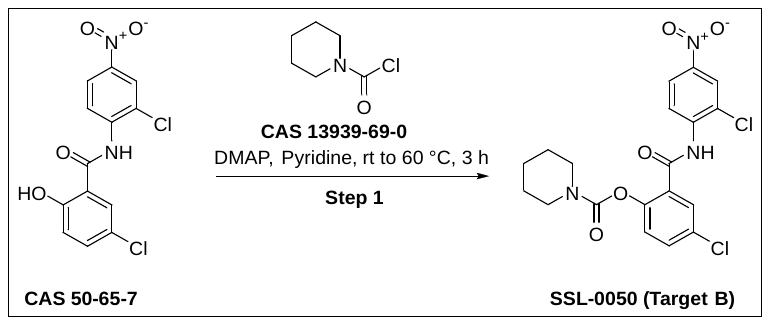

**Synthesis of 4-chloro-2-((2-chloro-4-nitrophenyl) carbamoyl) phenyl piperidine-1-carboxylate**

To a stirred solution of 5-chloro-N-(2-chloro-4-nitrophenyl)-2-hydroxybenzamide (CAS 50-65-7, 0.1 g, 0.3 mmol) in pyridine (2.0 mL) were added DMAP (CAS 1122-58-3, 0.022 g, 0.18 mmol) and piperidine-1-carbonyl chloride (CAS 13939-69-0, 0.103 g, 0.70 mmol) at rt and stirred at 60 °C for 3 h. The reaction mixture was diluted with water (30 mL) and extracted with EtOAc (3 x 15 mL). The combined organic layers were washed with 0.1N HCl solution (30 mL), dried over Na_2_SO_4_, filtered and concentrated under reduced pressure to afford 4-chloro-2-((2-chloro-4-nitrophenyl) carbamoyl) phenyl piperidine-1-carboxylate as an off-white solid (0.045 g, 0.10 mmol, 75% yield). ¹H NMR (400 MHz, DMSO-*d*_6_): *δ* 10.40 (s, 1H), 8.41 (d, *J* = 2.8 Hz, 1H), 8.29 (dd, *J* = 2.8, 2.4 Hz, 1H), 8.10 (d, *J* = 8.8 Hz, 1H), 7.76 (d, *J* = 2.4 Hz, 1H), 7.65 (dd, *J* = 2.4, 24 Hz, 1H), 7.35 (d, *J* = 8.8 Hz, 1H), 3.51 (brs, 2H), 3.35 (s, 2H), 1.53 - 1.44 (m, 2H), 1.44 - 1.45 (m, 4H), LCMS: O2h_LCMS_Method_A, Rt: 2.672 min, [M-H]^+^: 436.04.

**Experimental of SSL-0052 (APX-X-0011A)**

**Synthesis of 4-chloro-2-((2-chloro-4-nitrophenyl) carbamoyl) phenyl isopropyl carbonate**

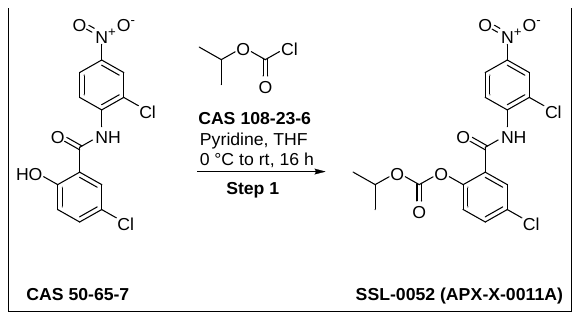

**Synthesis of 4-chloro-2-((2-chloro-4-nitrophenyl) carbamoyl) phenyl isopropyl carbonate**

To a stirred solution of 5-chloro-N-(2-chloro-4-nitrophenyl)-2-hydroxybenzamide (CAS 50-65-7, 0.5 g, 1.533 mmol) in THF (15.0 mL) were added Pyridine (0.133 g, 1.68 mmol) and isopropyl chloroformate (CAS 108-23-6; 0.376 g, 3.06 mmol) at 0 °C and stirred at rt for 16 h. The reaction mixture was diluted with water (100 mL) and extracted with EtOAc (2 x 100 mL). The combined organic layers were dried over Na_2_SO_4_, filtered and concentrated under reduced pressure. The crude material was purified by flash chromatography (Biotage, Normal phase, 230 - 400 mesh silica, 3% EtOAc in n-hexane) to afford 4-chloro-2-((2-chloro-4-nitrophenyl) carbamoyl) phenyl isopropyl carbonate as an off-white solid (0.135 g, 0.327 mmol, 21% yield). ¹H NMR (400 MHz, CDCl_3_): *δ* 9.55 (s, 1H), 8.89 (d, *J* = 9.2 Hz, 1H), 8.37 (d, *J* = 2.4 Hz, 1H), 8.24 (dd, *J* = 2.4, 2.4 Hz, 1H), 8.15 (d, *J* = 2.8 Hz, 1H), 7.58 (dd, *J* = 2.8, 2.4 Hz, 1H), 7.35 (d, *J* = 8.8 Hz, 1H), 5.05 - 4.99 (m, 1H), 1.38 (d, *J* = 6.4 Hz, 6H) LCMS: O2h_LCMS_Method_A, Rt: 2.672 min, [M-H]^+^: 411.0.

**Experimental of SSL-0053 (APX-X-0011B)**

**Synthesis of 4-chloro-2-((2-chloro-4-nitrophenyl) carbamoyl) phenyl isobutyl carbonate**

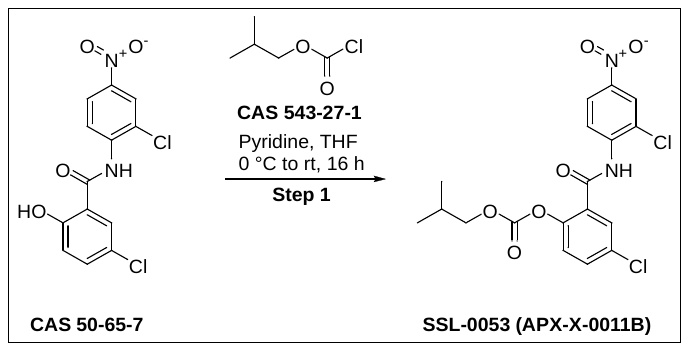

**Synthesis of 4****-chloro-2-((2-chloro-4-nitrophenyl) carbamoyl) phenyl isobutyl carbonate**

To a stirred solution of 5-chloro-N-(2-chloro-4-nitrophenyl)-2-hydroxybenzamide (CAS 50-65-7, 0.3 g, 0.92 mmol) in THF (15.0 mL) were added pyridine (0.08 g, 1.01 mmol) and isobutyl chloroformate (CAS 543-27-1, 0.138 g, 1.01 mmol) at 0 °C and stirred at rt for 16 h. The reaction mixture was diluted with water (50 mL) and extracted with EtOAc (2 x 50 mL). The combined organic layers dried over Na_2_SO_4_ filtered and concentrated under reduced pressure. The crude material was purified by flash chromatography (Biotage, Normal phase, 230 – 400 mesh silica, 1% EtOAc in hexane) afford 4-chloro-2-((2-chloro-4-nitrophenyl) carbamoyl) phenyl isobutyl carbonate as an off-white solid (0.15 g, 0.364 mmol, 39% yield). ¹H NMR (400 MHz, CDCl_3_): δ 9.51 (bs, 1H), 8.88 (d, *J* = 9.2 Hz, 1H), 8.37 (d, *J* = 2.8 Hz*,* 1H), 8.24 (dd, *J* = 2.4, 2.4 Hz, 1H), 8.13 (d, *J* = 2.8 Hz, 1H), 7.59 (dd, *J* = 2.8, 2.8 Hz, 1H), 7.35 (d, *J* = 8.8 Hz*,* 1H), 4.10 (d, *J* = 6.40 Hz*,* 2H), 2.08 – 2.01 (m *,* 1H), 0.98 (d, *J* = 6.8 Hz*,* 6H) LCMS: O2h_LCMS_Method_A, Rt: 2.851 min, [M-H]^+^: 425.1.

**Experimental of SSL-0058 (APX-X-0037)**

**Synthesis of** **methyl 3-(4-chloro-2-((2-chloro-4-nitrophenyl) carbamoyl) phenoxy)-2,2-dimethylpropanoate**

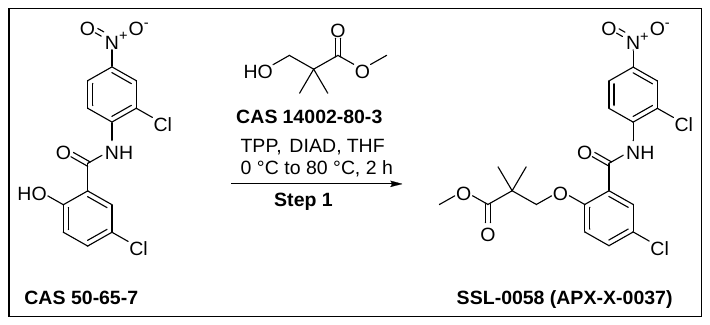

**Synthesis of methyl 3-(4-chloro-2-((2-chloro-4-nitrophenyl) carbamoyl) phenoxy)-2,2-dimethylpropanoate**

To a stirred solution of 5-chloro-N-(2-chloro-4-nitrophenyl)-2-hydroxybenzamide (CAS 50-65-7, 0.7 g, 2.147 mmol) and (CAS 14002-80-3, 0.425 g, 3.220 mmol) in THF (14.0 mL) was added TPP (1.125 g, 4.294 mmol) at 0 °C. After 5 min, DIAD (0.867 g, 4.294 mmol) was added dropwise and stirred at 80 °C for 2 h. The reaction mixture was diluted with water (20 mL) and extracted with EtOAc (2 x 50 mL). The combined organic layers were dried over Na_2_SO_4_, filtered and concentrated under reduced pressure. The crude material was purified by flash chromatography (Biotage, Normal phase, Silica gel, Mesh size – 230 - 400, 15% EtOAc in Hexane) followed by triturated with n-pentane (10.0 mL) to afford methyl 3-(4-chloro-2-((2-chloro-4-nitrophenyl) carbamoyl) phenoxy)-2,2-dimethylpropanoate as an off-white solid (0.1 g, 0.22 mmol, 10.0% yield). ¹H NMR (400 MHz, DMSO-*d*_6_): *δ* 10.08 (s, 1H), 8.43 (d, *J* = 1.2 Hz, 1H), 8.29 (d, *J =* 1.2 Hz, 2H), 7.71 (d, *J* = 2.8 Hz 1H), 7.61 (dd, *J* = 2.8, 2.8 Hz, 1H), 7.32 (d, *J =* 8.8 Hz*,* 1H), 4.21 (s, 2H), 3.47 (s, 3H), 1.22 (s, 6H), LCMS: O2h_LCMS_Method_A, Rt: 2.804 min, [M+H]^+^: 441.2.

**Experimental of SSL-0061 (APX-X-0013_By Product)**

**Synthesis of 4-chloro-2-((2-chloro-4-nitrophenyl) carbamoyl) phenyl butyrate**

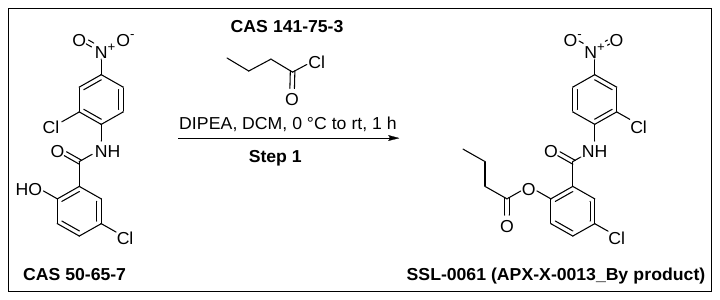

**Synthesis of 4-chloro-2-((2-chloro-4-nitrophenyl) carbamoyl) phenyl butyrate**

To a stirred solution of 5-chloro-N-(2-chloro-4-nitrophenyl)-2-hydroxybenzamide (CAS 50-65-7, 0.2 g, 0.613 mmol) in DCM (4.0 mL) were added DIPEA (0.237 g, 1.84 mmol) and butyryl chloride (CAS 141-75-3, 0.078 g, 0.736 mmol) at 0 °C and stirred at rt for 1 h. The reaction mixture was diluted with water (30 mL) and extracted with DCM (2 x 30 mL). The combined organic layers dried over Na_2_SO_4,_ filtered and concentrated under reduced pressure. The crude material was purified by trituration using MeOH (10 mL) to afford 4-chloro-2-((2-chloro-4-nitrophenyl) carbamoyl) phenyl butyrate as an off-white solid (0.08 g, 0.201 mmol, 33% yield). ¹H NMR (400 MHz, DMSO-*d*_6_): δ 10.49 (s, 1H), 8.41 (d, *J* = 2.4 Hz, 1H), 8.28 (dd, *J* = 2.8, 2.4 Hz, 1H), 8.08 (d, *J* = 9.2 Hz ,1H), 7.83 (d, *J* = 2.4 Hz ,1H), 7.70 (dd, *J* = 2.8, 2.4 Hz*,*1H), 7.35 (d, *J* = 8.8 Hz*,* 1H), 2.55 - 2.50 (m, 2H), 1.65 - 1.55 (m *,* 2H), 0.89 (t, *J* = 7.2 Hz, 3H), LCMS: O2h_LCMS_Method_A, Rt: 2.71 min, [M-H]^+^: 395.0

**3. Assessment of Thermodynamic Aqueous Solubility of Test Compound in Aqueous Buffer Systems.**

**Materials and Reagents:**

- Test compound (purity ≥ 98%)
- 10mM PBS pH 7.4.
- Syringe filter 13mm
- Syringe
- 1N Hydrochloric acid (HCl)
- 1N Sodium hydroxide (NaOH)
- Milli-Q water (HPLC grade)
- Dimethyl sulfoxide (DMSO, analytical grade)
- Microcentrifuge tubes
- Manual Pipettes
- 96 well uv plates
- Uv-visible spectrophotometer
- 50 ml concial centrifuge tubes

**Instrumentation:**

- Epoch-Biotek, uv-vis spectrophotometer
- Vortex
- pH meter (calibrated)
- Rotospin
- Sonicator
- Dry bath

**Methodology:**

I. **Preparation of Stock Solutions:** A 50 mM stock solution of the test compound was prepared in DMSO. If any precipitation was observed in preparation of main stock, sonicator was used.
II. **Preparation of aqueous Solutions:** ~ 1.0 mg of compound in to 1 mL of 10mM PBS buffer in an Eppendorf micro centrifuge tube was added and sealed with parafilm to avoid any leakage.
III. **Incubation Conditions:** The test compound microcentrifuge tubes were incubated at RT for 24 hours.
IV. **λmax:** Working stock of 1mM was prepared. Review the spectrophotometer data and identify the peak of the absorption. Prepare the linearity protocol and perform linearity using UV visible spectrophotometer.
V. **Sample Processing:** After 24 hours of incubation period the buffer solution in micro centrifuge tubes were filtered and aqueous concentration of the compound which is soluble in buffer was calculated.
VI. **Analytical Method:** Quantitative analysis was performed using uv-visible spectrophotometer.
VII. **Data Analysis:** The extent of test compound dissolution was determined in aqueous buffer system and was represented by plotting concentration vs OD line graph. Solubility of compound was determined using formula y=mx+c.
4. **Assessment of Chemical Stability of Test Compound in Aqueous Buffer Systems**

Objective: To evaluate the chemical stability of the test compound under physiologically relevant pH conditions using a validated buffer-based method.

Materials and Reagents:

· Test compound (purity ≥ 98%)

· Potassium dihydrogen phosphate (KH₂PO₄)

· Dipotassium hydrogen phosphate (K₂HPO₄)

· 1N Hydrochloric acid (HCl)

· 1N Sodium hydroxide (NaOH)

· Milli-Q water (HPLC grade)

· Dimethyl sulfoxide (DMSO, analytical grade)

· Phosphate buffer solutions prepared at pH 2.0, 4.0, 6.8, and 8.0

· LC-MS grade acetonitrile or methanol

Instrumentation:

· HPLC-MS/MS system equipped with C18 reverse-phase column

· Centrifuge capable of 10,000 rpm

· pH meter (calibrated)

· Water bath or incubator (maintained at 37 ± 1 °C)

Methodology:

I. Preparation of Stock Solutions: A 10 mM stock solution of the test compound was prepared in DMSO. Working solutions were diluted to a final concentration of 10 µM in 100 mM phosphate buffer at the desired pH (2.0, 4.0, 6.8, 8.0). The final DMSO concentration in all samples was kept ≤1% (v/v).
II. Incubation Conditions: The buffered test solutions were incubated in sealed polypropylene tubes at 37 ± 1 °C. Aliquots were withdrawn at predetermined time points: 0, 0.5, 1, 2, 4, 8, and 24 hours.
III. Quenching: At each time point, an aliquot was transferred to a pre-cooled microcentrifuge tube and immediately mixed with equal volume of ice-cold acetonitrile (containing an internal standard, if applicable) to stop the reaction.
IV. Sample Processing: Samples were vortexed and centrifuged at 10,000 rpm for 10 minutes. The supernatant was transferred to HPLC vials for analysis.
V. Analytical Method: Quantitative analysis was performed using HPLC-MS/MS. The analyte was separated on a C18 reverse-phase column under gradient conditions using water and acetonitrile containing 0.1% formic acid. The mass transition for the test compound was monitored in multiple reaction monitoring (MRM) mode.
VI. Data Analysis: The peak area ratio of the test compound (relative to internal standard) was determined at each time point. The percentage remaining was calculated by comparing the peak area at each time point to the time zero sample. A semi-log plot of % remaining versus time was generated to estimate the degradation kinetics.

Interpretation: The stability profile was assessed across different pH levels to understand the susceptibility of the compound to hydrolytic degradation under physiologically relevant conditions. Compounds showing ≥85% remaining at 24 hours were considered chemically stable.

5. **Assessment of Kinetic Aqueous Solubility of Test Compound in Aqueous Buffer Systems**

Objective: To determine the aqueous solubility of the test compound under physiologically relevant pH conditions using a validated and standardized kinetic aqueous assay.

Materials and Reagents:

· Test compound (purity ≥ 98%)

· 10mM PBS pH 7.4.

· Glass tubes

· Syringe filter 13mm

· Syringe

· 1N Hydrochloric acid (HCl)

· 1N Sodium hydroxide (NaOH)

· Milli-Q water (HPLC grade)

· Dimethyl sulfoxide (DMSO, analytical grade)

· Microcentrifuge tubes

· Manual Pipettes

· 96 well uv plates

· Uv-visible spectrophotometer

· 50 ml concial centrifuge tubes

Instrumentation:

· Epoch-Biotek, uv-vis spectrophotometer

· Vortex

· pH meter (calibrated)

· Rotospin

· Sonicator

· Dry bath

Methodology:

I. Preparation of Stock Solutions: A 50 mM stock solution of the test compound was prepared in DMSO. If any precipitation was observed in preparation of main stock, sonicator was used. Test compound was spiked in 1ml of 10mM PBS buffer starting from 0.5%, 1%, 1.5% and 2% which means 5μL, 10μL, 15μL and 20µl of compound respectively. Spiking was halted immediately after precipitation was observed.
II. Incubation Conditions: The buffered test solutions were incubated in sealed glass tubes at RT for 4 hours.
III. λmax: Working stock of 1mM was prepared. Review the spectrophotometer data and identify the peak of the absorption. Prepare the linearity protocol and perform linearity using UV visible spectrophotometer.
IV. Sample Processing: After 4 hours of incubation period the buffer solution in glass tubes were filtered and aqueous concentration of the compound which is soluble in buffer was calculated.
V. Analytical Method: Quantitative analysis was performed using uv-visible spectrophotometer.
VI. Data Analysis: The extent of test compound dissolution was determined in aqueous buffer system and was represented by plotting concentration vs OD line graph. Solubility of compound was determined using formula y=mx+c.

Interpretation: The solubility profile was assessed using a validated buffer system. Compounds with <10 µM solubility were classified as low solubility, those with solubility between 10–100 µM were considered moderately soluble, and compounds exhibiting >100 µM solubility were categorized as highly soluble.

6. **Assessment of plasma Stability of Test Compound**

Objective: To assess the stability of the test compound in human plasma under physiologically relevant conditions using a validated and standardized plasma stability assay.

Materials and Reagents:

· Test compound (purity ≥ 98%)

· 1N Hydrochloric acid (HCl)

· 1N Sodium hydroxide (NaOH)

· Milli-Q water (HPLC grade)

· Dimethyl sulfoxide (DMSO, analytical grade)

· LC-MS grade acetonitrile or methanol

· Human Plasma (K2EDTA)

Instrumentation:

· HPLC-MS/MS system equipped with C18 reverse-phase column

· Centrifuge capable of 10,000 rpm

· Vortex

· pH meter (calibrated)

· Water bath or incubator (maintained at 37 ± 1 °C)

Methodology:

I. Preparation of Stock Solutions: A 50 mM stock solution of the test compound was prepared in DMSO. Working solutions were diluted to a final concentration of 10 µM in 100% plasma. The final DMSO concentration in all samples was kept 2% (v/v).
II. Incubation Conditions: The master mix containing plasma with test solutions were incubated in sealed polypropylene tubes at 37 ± 1 °C. Aliquots were withdrawn at predetermined time points: 0, 15, 30, 60, 90 and 120 mins.
III. Quenching: At each time point, an aliquot was transferred to a pre-cooled microcentrifuge tube and immediately mixed with 1:4 volume of ice-cold acetonitrile (containing an internal standard, if applicable) to stop the reaction.
IV. Sample Processing: Samples were vortexed and centrifuged at 10,000 rpm for 10 minutes. The supernatant was transferred to HPLC vials for analysis.
V. Analytical Method: Quantitative analysis was performed using HPLC-MS/MS. The analyte was separated on a C18 reverse-phase column under gradient conditions using water and acetonitrile containing 0.1% formic acid. The mass transition for the test compound was monitored in multiple reaction monitoring (MRM) mode.
VI. Data Analysis: The peak area ratio of the test compound (relative to internal standard) was determined at each time point. The percentage remaining was calculated by comparing the peak area at each time point to the time zero sample. A semi-log plot of % remaining versus time was generated to estimate the degradation kinetics.

Interpretation: The stability profile was evaluated in human plasma to assess the compound's degradation under physiologically relevant conditions. Compounds retaining ≥90% of the parent form after 2 hours were classified as stable, those with 60–90% remaining were considered moderately stable, and compounds showing <60% remaining within 1–2 hours were categorized as unstable.

Fig. S1 **SSL0024 and SSL-0061 show best HCC inhibition efficacy**. Huh7, HepG2, and Hep3B cells were treated with compounds for 72 hours. Data are presented as Mean ± SEM. n = 3.

Fig. S2 **Tissue niclosamide concentrations derived from additional screened prodrug candidates**. The oral dose is 100 mg/kg for all compounds. SSL-0024, SSL-0052, SSL-0053 treated tissues were harvested after 4 hours of the administration, the rest of two were harvested at 24-hour time point. Compounds were administered by oral gavage (n = 5 per group). Data are presented as Mean ± SEM. n =6, 5, 4, 5, 5 in SSL-0024, SSL-0052, SSL-0053, SSL-0058, and SSL-0058 group, respectively.

Table S1 **Liver, kidney function, and lipid metabolism assay**. Blood was collected after 24 h of compound administration. Data are presented as mean ± SEM. n = 5. **p* < 0.05, ***p* < 0.01, ****p* < 0.001 vs. control.

WBC white blood cell, RBC red blood cell, HGB hemoglobin, HCT Hematocrit, MCV mean corpuscular volume, MCH mean corpuscular hemoglobin, MCHC mean corpuscular hemoglobin concentration, RDW red cell distribution width, PDW Platelet distribution width, MPV mean platelet volume, p-lcr Platelet-large cell ratio, PCT Procalcitonin, IRF immature reticulocyte fraction, IFR immature reticulocyte fraction, MFR medium fluorescence reticulocytes, AST Aspartate Transferase, ALT Alanine Aminotransferase, GGT Gamma-glutamyl Transferase , BUN Blood urea nitrogen. NA: Not available
